# A *Mesp1*-dependent developmental breakpoint in transcriptional and epigenomic specification of early cardiac precursors

**DOI:** 10.1101/2022.08.22.504863

**Authors:** Alexis Leigh Krup, Sarah A.B. Winchester, Sanjeev S. Ranade, Ayushi Agrawal, W. Patrick Devine, Tanvi Sinha, Krishna Choudhary, Martin H. Dominguez, Reuben Thomas, Brian L. Black, Deepak Srivastava, Benoit G. Bruneau

## Abstract

Transcriptional networks governing cardiac precursor cell (CPC) specification are incompletely understood due in part to limitations in distinguishing CPCs from non-cardiac mesoderm in early gastrulation. We leveraged detection of early cardiac lineage transgenes within a granular single cell transcriptomic time course of mouse embryos to identify emerging CPCs and describe their transcriptional profiles. *Mesp1*, a transiently-expressed mesodermal transcription factor (TF), is canonically described as an early regulator of cardiac specification. However, we observed perdurance of CPC transgene-expressing cells in *Mesp1* mutants, albeit mis-localized, prompting us to investigate the scope of *Mesp1*’s role in CPC emergence and differentiation. *Mesp1* mutant CPCs failed to robustly activate markers of cardiomyocyte maturity and critical cardiac TFs, yet they exhibited transcriptional profiles resembling cardiac mesoderm progressing towards cardiomyocyte fates. Single cell chromatin accessibility analysis defined a *Mesp1*-dependent developmental breakpoint in cardiac lineage progression at a shift from mesendoderm transcriptional networks to those necessary for cardiac patterning and morphogenesis. These results reveal *Mesp1*-independent aspects of early CPC specification and underscore a *Mesp1-*dependent regulatory landscape required for progression through cardiogenesis.

## INTRODUCTION

Cardiogenesis requires precise specification and patterning of the cardiac precursor cells (CPCs) as they emerge from the gastrulating mesoderm in very early stages of embryogenesis. Errors in this process lead to congenital heart defects (CHDs), which affect 1-2% of live births (Bruneau, 2008). The genetic etiology of CHDs indicates that genes encoding transcriptional regulators are overrepresented as causative and are predominantly haploinsufficient, indicating that fine dysregulation of gene expression is a critical mechanism for disease (Zug, 2022; Nees & Chung, 2019). A thorough delineation of the transcriptional networks governing cardiogenesis is foundational to understanding how defects in this process manifest as CHDs, and may inform the design of strategies to treat CHDs and heart disease broadly.

Cardiogenesis begins when mesoderm progenitors emerge from the primitive streak and migrate towards the anterior-lateral aspects of the developing embryo (Saga, Kitajima & Miyagawa-Tomita, 2000; Saga et al., 1999). Interrogating the earliest cardiac progenitors distinctly from the developing mesoderm has historically been challenging due to a paucity of molecular markers available to distinguish a CPC from the rest of the developing mesoderm. Prior studies used lineage tracing of mesoderm progenitors expressing the basic-helix-loop-helix (bHLH) transcription factor (TF) *Mesp1,* which is transiently expressed in cells that go on to contribute to the heart, somitic mesoderm derivatives, and craniofacial mesoderm (Devine et al., 2014; Lescroart et al., 2014; Saga et al., 1999). Clonal lineage tracing studies have shown that a subset of *Mesp1*+ cells at early gastrulation are fated for distinct cardiac substructures well before anatomy is patterned, highlighting extensive diversification among early mesodermal progenitors (Devine et al., 2014; Lescroart et al., 2014; Liu, 2017).

Deletion of *Mesp1* in mice variably disrupts specification and migration of cardiac progenitors (Ajima et al., 2021; Saga, Kitajima & Miyagawa-Tomita, 2000; Saga et al., 1999; Kitajima et al., 2000; Lescroart et al., 2018). During in vitro cardiac differentiation, overexpression of *Mesp1* induces expression of subsequent cardiac TFs, indicating a potentially instructive role in cardiogenesis (Chiapparo et al., 2016; Bondue et al., 2008; Lindsley et al., 2008; Wu, 2008; Kelly, 2016; Bondue & Blanpain, 2010; Lin et al., 2022; Soibam et al., 2015). Gain of function experiments suggest a broad and important function for *Mesp1* in mesoderm differentiation, but the *in vivo* gene regulatory landscape controlled by *Mesp1* remains unclear (Costello et al., 2011; Saga et al., 1999; Liu, 2017; Saga, Kitajima & Miyagawa-Tomita, 2000; Ajima et al., 2021; Saga et al., 1996; Kitajima et al., 2000; Lin et al., 2022).

Previous studies identified an enhancer of *Smarcd3,* “F6”, which is specifically active in CPCs fated to become the totality of heart cells, and is active shortly after *Mesp1* expression and before other early cardiac-specific TFs are expressed (Devine et al., 2014; Yuan et al., 2018). Thus, *Smarcd3*-F6 activity enables distinct identification of CPCs as they emerge from the developing mesoderm. We found that the *Smarcd3*-F6 enhancer remains active in posterior regions of *Mesp1* KOs, indicating perdurance of cardiogenesis in some capacity. Here, we utilized the *Smarcd3*-F6 transgene to comprehensively delineate the dynamic transcriptional and epigenomic consequences of *Mesp1* loss during early cardiogenesis and reveal *Mesp1*-independent aspects of cardiac specification. This study challenges the concept of a master regulator for cardiac specification by defining transcriptional phases with different vulnerabilities to *Mesp1* loss.

## RESULTS

### Computational detection of fluorescent transgene reporters enables identification of emerging cardiogenic mesoderm from whole embryo single cell transcriptomic data

To identify the emerging cardiogenic mesoderm cells at their earliest stages we employed a reporter transgene strategy in combination with scRNA-seq on a whole embryo time course spanning early gastrulation (E6.0) until cardiac crescent stages (E7.75) (Fig. 1A-B, Fig. S1). Embryos contained a fluorescent transgene reporter for the *Mesp1* lineage via *Mesp1^Cre^;Rosa26R^Ai14^* (Saga et al., 1999; Madisen et al., 2010), and the *Smarcd3*-F6::eGFP enhancer transgene that constitutively marks CPCs (Devine et al., 2014; Yuan et al., 2018) (Fig. 1C, Fig. S1). We processed whole embryos for scRNA-seq and used computational detection of the fluorescent transgene reporters to identify the emerging cardiogenic mesoderm (Fig. 1C). Following an annotation of cell types in the whole embryo atlas (Fig. S2A, Table S1, Table S2A), we subsetted cell clusters expressing the Ai14 and eGFP fluorescent transgenes and demonstrated that these cells resemble the emerging cardiac mesoderm (Fig. 1D-F, Fig. S2B, Table S1, Table S2B). *Smarcd3*-F6+ cell clusters (Fig. 1F, Fig. 2B, Table S1, Table S2B) co-expressed early cardiac and mesoderm genes in the mesoderm exiting the primitive streak (meso exiting PS), anterior mesendoderm (antME), and cells of the lateral plate mesoderm (LPM) such as *T, Eomes*, *Mesp1*, *Mixl1,* and *Smarcd3* (Fig. 1D, Fig. S3A). *Smarcd3*-F6+ cardiomyocytes (CMs) co-expressed cardiac structural genes such as *Myl7*, *Tnnt2*, *Actc1* (Fig. 1D-F, Fig. S3B) and cardiac TFs such as *Tbx5*, *Hand1*, *Nkx2-5, Gata5* (Fig. 1D-F, Fig. S3C). Thus, *Smarcd3*-F6 enhancer transgene expressing cells have transcriptional signatures of early emerging CPCs, extending the initial description of this transgene (Devine et al., 2014) by validating high-fidelity demarcation of emerging early CPCs within the mesoderm prior to expression of cardiac-specific TFs in scRNA-seq data (Fig. S2B, Fig. S3).

**Fig. 1.**
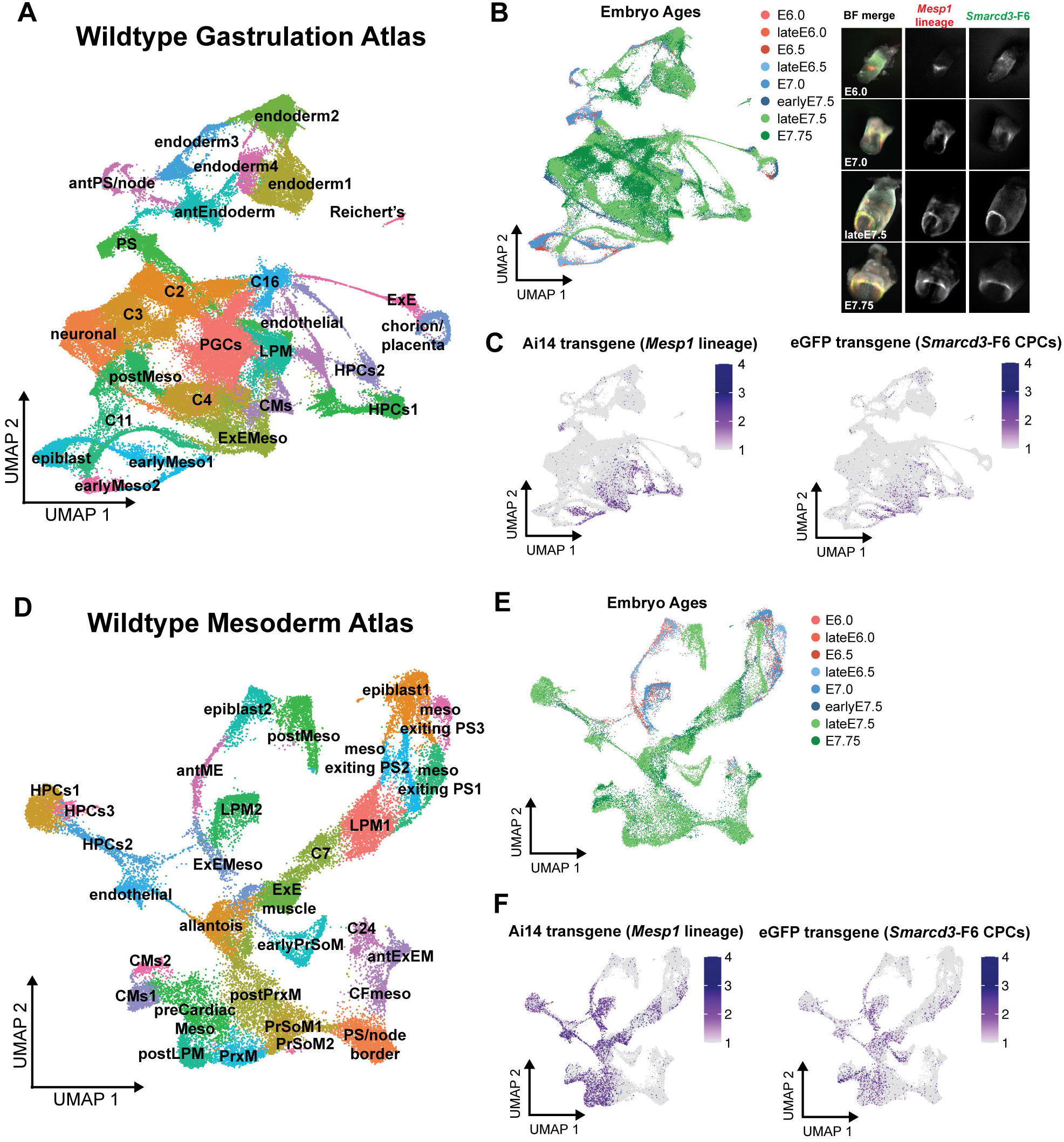
Identification of the emerging cardiogenic mesoderm using fluorescent transgenes in whole embryo single cell transcriptomic data. (A) Uniform manifold approximation and projection (UMAP) of 94,824 cells representing 27 cell types from gastrulating embryos. (B) UMAP labeled with embryo ages included in atlas and representative embryo images showing domains of fluorescent Ai14 (*Mesp1* lineage) and eGFP (*Smarcd3*-F6) transgenes. Images not scaled. (C) UMAP feature plots showing expression of fluorescent transgenes isolated to mesodermal cell types. (D) UMAP of 34,724 mesodermal cells subsetted from full atlas, representing 30 cell types. (E) UMAP labeled with embryo ages and (F) UMAP feature plots showing expression of fluorescent transgenes, Ai14 for *Mesp1* lineage and eGFP for *Smarcd3*-F6+ CPCs.

**Fig. 2.**
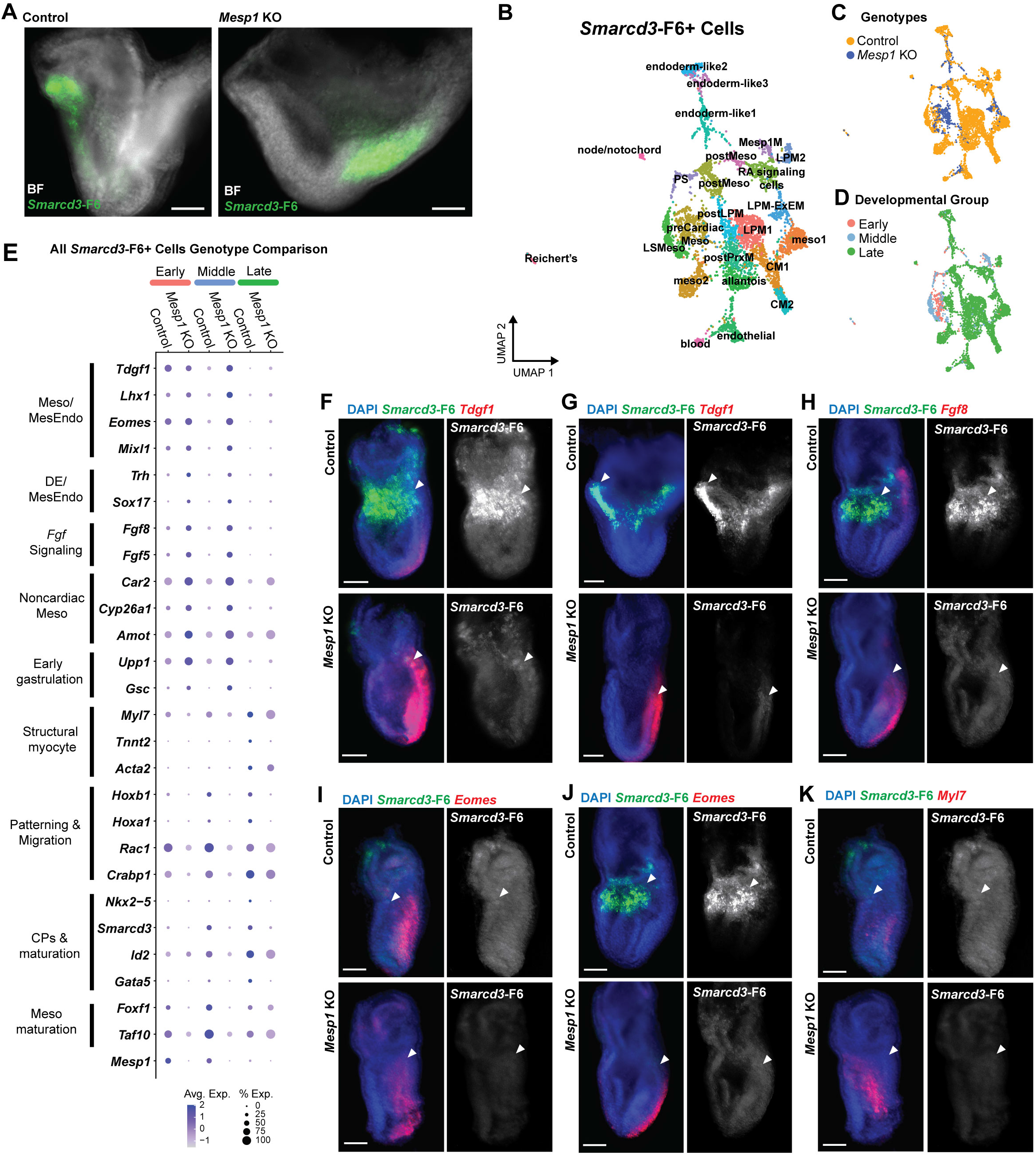
Transcriptional profiles of *Smarcd3*-F6+ cells in *Mesp1* KO embryos. (A) Fluorescence *in situ* hybridization for *Smarcd3*-F6 expression (green) in cardiac crescent stage (E7.75) *Mesp1* KO and control littermate embryos. (B) UMAP atlas of 4,868 *Smarcd3*-F6+ cells representing 24 cell types. (C-D) UMAPs colored by (C) genotype and (D) relative developmental stages, Early (E6.0-E6.5), Middle (late E6.5 – E7.5), Late (late E7.5-early E7.75). (E) Dotplot representation of gene expression across genotypes at relative developmental stages. Size of dot denotes percent of cells expressing gene, color of dot represents average gene expression. (F-K) Multiplexed fluorescence *in situ* hybridization for *Smarcd3*-F6 (green) and (F-G) *Tdgf1* (red) in representative (F) Early and (G) Middle stages, (H) *Fgf8* (red) in Middle stages, (I-J) *Eomes* (red) in (I) Early and (J) Middle stages, (K) *Myl7* (red) in Early stages. Arrowheads denote *Smarcd3*-F6+ cardiogenic regions. Scale bars are 100 μm.

In summary, we generated a resource dataset and interrogated dynamic gene expression programs in the cardiogenic mesoderm using reporter transgenes, which will facilitate description of emerging heterogeneity within the cardiac lineage during gastrulation.

### Transcriptional profiling of *Smarcd3*-F6+ cells shows enduring expression of cardiac genes in *Mesp1* knockout embryos

To determine the requirement for *Mesp1* in establishing CPC identity, we investigated the transcriptional identities of *Smarcd3*-F6+ cells upon loss of *Mesp1*. We detected *Smarcd3*-F6 expressing cells in *Mesp1^Cre/Cre^* (*Mesp1* KO) embryos, although positive cells are localized posteriorly relative to control embryos at early cardiac crescent stages (Fig. 2A). The persistence of *Smarcd3*-F6+ cells led us to hypothesize that these cells represent retained CPCs, suggesting that as previously described (Saga, Kitajima & Miyagawa-Tomita, 2000; Ajima et al., 2021; Saga et al., 1999), aspects of early cardiac specification may be *Mesp1-*independent. Thus, the transcriptional and epigenomic programs regulated by and independent of *Mesp1* remain to be understood during *in vivo* cardiogenesis.

We performed scRNA-seq on whole *Mesp1* KO embryos and littermate controls along a timeline of developmental stages for early cardiogenesis spanning early gastrulation (E6.0) to cardiac crescent formation (E7.75) (Fig. S6A). We bioinformatically identified *Smarcd3*-F6-eGFP-expressing cells from the whole embryo time course (Fig. S6A-D, Table S4A) to generate an atlas of 4,868 *Smarcd3*-F6+ cells representing 24 cell types (Fig. 2B, Fig. S4A, Table S1, Table S3A). The majority of *Smarcd3*-F6+ cells represented early cardiac mesodermal derivatives such as the late streak mesoderm (LSMeso), *Mesp1*+ mesoderm (Mesp1M), posterior mesoderm (postMeso), LPM, precardiac mesoderm (preCardiacMeso), and early CMs (Fig. 2B-C, Fig. S4A). We detected cells of the allantois, lateral plate mesoderm/extraembryonic mesoderm (LPM-ExEM), and the node/notochord, consistent with *Smarcd3* expression in these domains (Fig 2B, Fig. S4B) (Takeuchi et al., 2007; Devine et al., 2014). Additionally, we found populations of blood, endothelial cells, Reichert’s membrane, posterior paraxial mesoderm (postPrxM) and cells appearing endoderm-like, potentially representing early mesendoderm cells (Fig. 2B, Fig. S4A). Inducible lineage labeling of *Smarcd3*-F6+ cells at E6.5 excluded lineage contributions to non-cardiac cell types (Devine et al., 2014), suggesting detection here is the result of genotype-agnostic, weak, or transient transgene expression (Fig. S4A).

To examine overall trends in gene expression differences between *Mesp1* KO and control, we performed a comparison of all *Smarcd3*-F6+ cells between genotypes irrespective of cell type or embryonic stage in the developmental timeline (Fig. S4B, Table S3B). We found that *Mesp1* KO *Smarcd3*-F6+ cells express mesodermal genes of the emerging cardiac lineage such as *Tdgf1*, *Lhx1*, *Eomes*, and *Myl7*, however mostly lacked expression of more mature cardiac progenitor markers such as *Nkx2-5* (Fig. 2E, Fig. S4A). When we divided the “all cells” genotype analysis into relative developmental stages separating “Early” embryos (E6.0-E6.5), “Middle” embryos (late E6.5-E7.5), and “Late” embryos (late E7.5 to early E7.75), we found that genotype discrepancies in cardiac-related gene expression were minor at Early stages and diverged with increasing embryonic age (Fig. 2E).

Relatedly, the distribution of genotypes across *Smarcd3*-F6+ cell types shows that *Mesp1* KO cells are not fully represented in every cell type (Fig. 2B-D, Fig. S4C). Both genotypes were present in mesoderm clusters (C1, C3), the preCardiacMeso (C4), the postMeso (C5), retinoic acid signaling cells (C6), LSMeso (C7), allantois (C8), endothelial (C9), postPrxM (C10), the endoderm-like clusters (C11, C14, C19), the LPM-ExEM cluster (C15), the primitive streak (PS) (C17), postMeso (C20), blood (C21), and Reichert’s (C23) (Fig. 2B-C, Fig. S4C). Only control cells were present in LPMs (C0, C16), CMs (C2, C12), postLPM (C13), Mesp1M (C18), node/notochord (C22). Many of the cell types comprised only of control were Late-stage embryo cells (Fig. 2B-D, Fig. S4C), indicating that cell type heterogeneity was affected with loss of *Mesp1* in *Smarcd3*-F6+ cells with increasing severity as development progresses. Furthermore, while the preCardiacMeso and LSMeso cell types were represented by both genotypes in Early- and Middle-staged embryos, the Late-stage embryo cells represented in the preCardiacMeso were exclusively *Mesp1* KO (Fig. 2B-D, Fig. S4C), indicating retention of precursor transcriptional profiles.

To understand *Mesp1*-correlated differences in emerging *Smarcd3*-F6+ CPCs in individual cell types, we performed differential expression testing within cell types present in both genotypes (Table S3C-F). Within preCardiacMeso and LSMeso cells, we found similar expression of *Tdgf1*, *Eomes, Fgf8*, genes involved in early mesoderm specification (Fig. S5A-C) (Probst et al., 2020; Reifers et al., 2000). These results were confirmed by multiplexed RNA *in situ* hybridization, which showed co-expression of *Smarcd3*-F6 with these markers in cardiogenic regions of E6.0-E6.5 (Fig. 2F, Fig. 2I) and E7.0 (Fig. 2H) embryos. Notably, *Mesp1* KO embryos at early stages showed decreased or delayed expression of *Smarcd3*-F6 (Fig 2H-K), and broad posterior expansion of *Tdgf1* (Fig. 2F) and *Fgf8* (Fig. 2H) expression beyond *Smarcd3*-F6+ cardiogenic regions. Additionally, *Tdgf1* and *Eomes* expression aberrantly perdured through late E7.5 (Fig. 2F,G) and E7.0 (Fig. 2I-J), respectively. Other genes involved in early mesoderm specification (*Fgf10),* lineage specification and pluripotency exit (*Chchd2* and *Nme2*), and non-cardiac mesoderm genes (*Amot*) were upregulated in *Mesp1* KO cells relative to controls, while genes involved in migration and patterning (*Lefty2*, *Rac1*, *Foxf1*) were downregulated (Fig. S5B,C) (Zhu et al., 2009, 2016; Migeotte, Grego-Bessa & Anderson, 2011; Sang et al., 2021).

Within the Late-stage-dominated LPM-ExEM and endoderm-like1 cell types, we found similar expression levels of *Myl7* between genotypes (Fig. 2K, Fig. S5A, Fig. S5D,E). *Mesp1* KO cells displayed relative upregulation of early mesoderm specification genes (*Tdgf1, Eomes, Fgf8, S100a10, Ifitm2*, *Fn1)* and downregulation of morphogenesis and migration genes (*Dlk1, Elavl1*) (Fig. S5D,E) (Probst et al., 2020; Cheng et al., 2013; Klymiuk et al., 2012; Saykali et al., 2019; Katsanou et al., 2009)

Collectively, these analyses indicate that the *Mesp1* KO transcriptional phenotype of *Smarcd3*-F6+ cells becomes increasingly disrupted as embryonic development progresses, consistent with the divergent morphology of *Mesp1* KO embryos at cardiac crescent stages (Fig. 2A-D).

### Alterations to cardiac mesoderm in *Mesp1* KO embryos become increasingly severe as gastrulation progresses

Following characterization of *Mesp1* KO effects in *Smarcd3*-F6+ cells specifically, we sought to understand alterations to the mesoderm, inclusive of *Smarcd3*-F6+ cells and the cardiac mesoderm, more broadly. We applied our method of dual-reporter transgene identification (Fig. 1) to generate an atlas of 35,792 mesodermal cells from both control and *Mesp1* KO embryos (Fig. S6A-F, Table S1, Table S4). The relative Early- and Middle-stage embryos showed a similar census of mesodermal cell types between genotypes, including preCardiacMeso (Fig. S6E-I). However, *Mesp1* KO Late-stage embryo mesoderm lacked many of the cell types present in control, such as mature CMs, PrxM, and PrSoM cells (Fig. S6E-I). To interrogate how these changes occur in developmental time, we divided the mesoderm dataset into the Early, Middle, and Late developmental stages as defined in Fig. 2E. Mesodermal cells for each stage were re-clustered, and differential gene expression was assessed between genotypes (Fig. 3, Table S4).

**Fig. 3.**
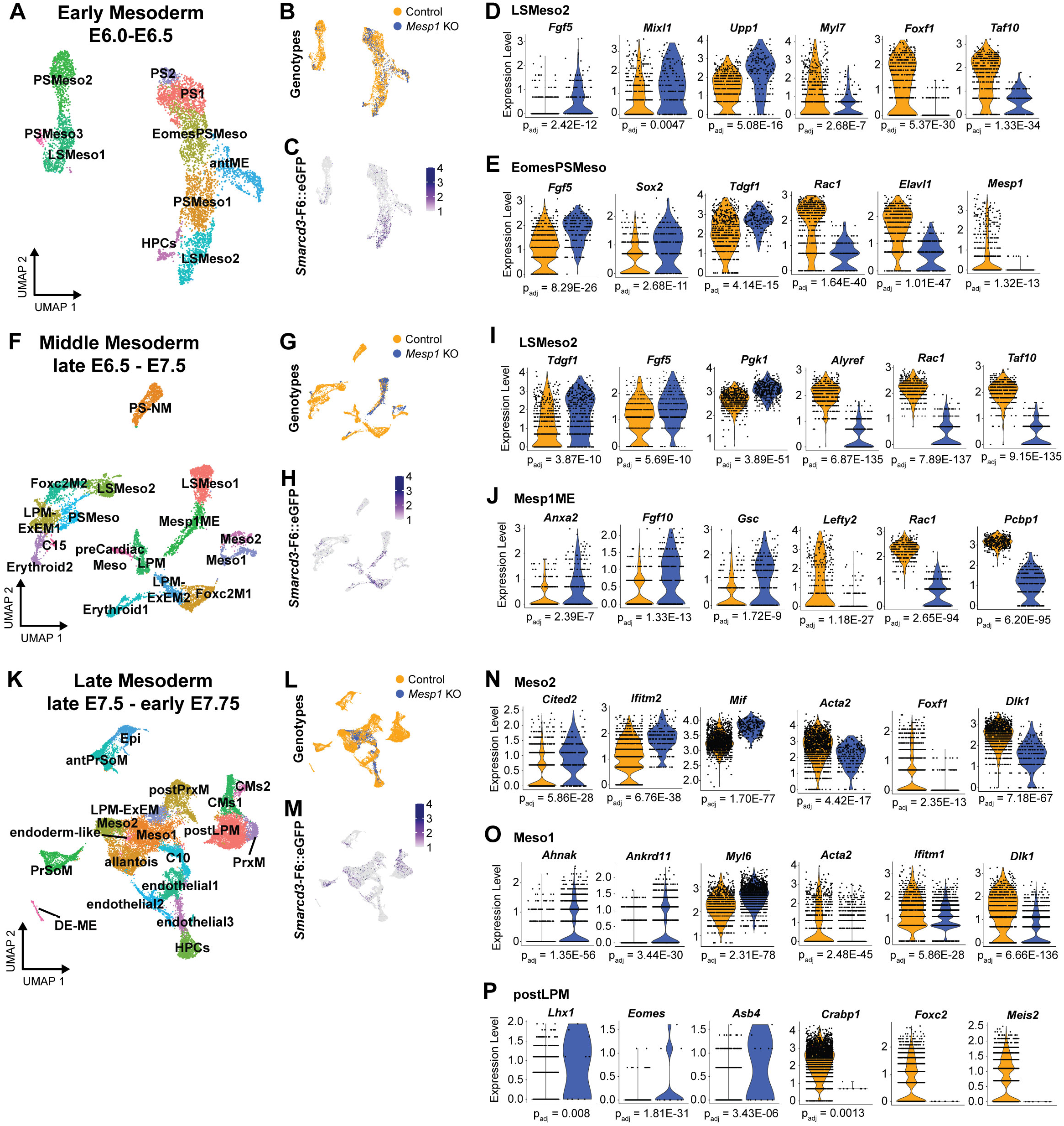
Transcriptional profiles of cardiac mesoderm in *Mesp1* KO embryos. Mesoderm scRNA-seq UMAP atlases for (A) Early (5,504 cells), (F) Middle (7,666 cells), and (K) Late (22,622 cells) developmental stages. Associated UMAPs for each stage atlas colored by (B, G, L) genotype and (C, H, M) *Smarcd3*-F6-eGFP expression. Differentially expressed genes in Early mesoderm in (D) LSMeso2 and (E) EomesPSMeso. Differentially expressed genes in Middle mesoderm in (I) LSMeso2and (J) Mesp1ME. Differentially expressed genes in Late mesoderm in (N) Meso2, (O) Meso1, and (P) postLPM. Significant changes are denoted with adjusted p values < 0.05.

Within the Early mesoderm dataset (Fig. 3A, Fig. S7A, Table S4C), we identified the LSMeso2 and *Eomes*+ primitive streak mesoderm (EomesPSMeso) as clusters of interest for cardiac specification based on enriched *Smarcd3*-F6+ expression (Fig. 3C). Both genotypes were present in each cell type (Fig. 3B), indicating that *Mesp1* KO cells are able to engage with transcriptional programs to exit pluripotency and initiate cardiac mesoderm specification. Differential gene expression analysis revealed *Mesp1* KO cells showed upregulation of mesendoderm and PS markers (*Fgf5*, *Mixl1*, *Upp1*, *Fgf5*, *Sox2*, *Tdgf1*), downregulation of LPM differentiation genes (*Foxf1*, *Taf10*), downregulation of migration and patterning genes (*Rac1*, *Elavl1*), and persistent but decreased expression of cardiac *Myl7* (Fig. 3D-E, Table S4D,E).

Within the Middle mesoderm dataset (Fig. 3F-H, Fig. S7B, Table S4F) we focused on the *Smarcd3*-F6 enriched *Mesp1*+ mesendoderm cluster (Mesp1ME) and its developmental predecessors, LSMeso2 cells. Middle-stage LSMeso2 cells (Fig. 3I, Table S4G) showed similar expression patterns between genotypes to Early-stage LSMeso cells (Fig. 3D). *Mesp1* KO cells of the Mesp1ME upregulated posterior mesoderm organization genes (*Fgf10* and *Gsc*) (Probst et al., 2020; Meijer et al., 2000; Branney et al., 2009) and the non-cardiac mesoderm gene *Anxa2* (Schwartz et al., 2014; Wang et al., 2015), and downregulated *Lefty2, Rac1*, and myogenesis differentiation gene *Pcbp1* (Shi & Grifone, 2021) (Fig. 3J, Table S4H). Notably, there was an absence of *Mesp1* KO cells in *Smarcd3*-F6 and *Mesp1* enriched clusters representing *Foxc2+* mesoderm cells (Fig. 3F-H, Fig. S8A). *Foxc2* operates in cardiac field diversification and morphogenesis (Seo & Kume, 2006; Lescroart et al., 2018). Examination in E6.75 embryos by immunohistochemistry and light sheet imaging showed that anterior-proximal marker domains were misaligned in *Mesp1* KO embryos, and *Foxc2* was absent (Fig. S8A-B). Together, these results indicate dysregulation of networks controlling cellular movements and domain boundaries, as well as reduced cellular diversification in *Mesp1* KO embryos of pre-crescent stages.

Analysis of Late mesoderm *Mesp1* KO cells revealed restricted diversity of both cardiac and other mesodermal cell types (Fig. 3K-L, Fig. S7C, Table S4I). Furthermore, while both genotypes were found in *Smarcd3*-F6 enriched clusters (Meso1, Meso_2) and the postLPM, there were no *Mesp1* KO cells in the CM clusters (Fig. 3K,M). *Mesp1* KO cells from Meso1 and Meso2 clusters had highly disrupted transcriptional profiles characterized by upregulation of several mesodermal genes (*Cited2*, *Ifitm2*, *Mif*, *Ahnak, Ankrd11, Myl6*) (Weninger et al., 2005; Lange et al., 2003; Huang et al., 2022) and downregulation of cardiac maturation genes (*Dlk1*, *Acta2, Ifitm1*) (Pursani et al., 2017; Klymiuk et al., 2012) (Fig. 3N,O, Table S4J,K). Additionally, the few *Mesp1* KO cells present in the postLPM cluster upregulated genes involved in mesendoderm specification and organization (*Lhx1, Eomes, Asb4*) (Fernandez-Guerrero et al., 2021) and downregulated or else lacked patterning, morphogenesis, and maturation genes (*Crabp1*, *Foxc2*, *Meis2*) (Fig. 3P, Table S4L). From these results we conclude that Late stage *Mesp1* KO embryos fail to produce mature CMs and various mesoderm cell types, and display highly disrupted transcriptional profiles in the cardiac mesoderm cells that are present.

Thus, similar to the patterns described specifically in *Mesp1* KO *Smarcd3*-F6+ CPCs, *Mesp1* KO cardiac mesoderm cells show transcriptional dysregulation that becomes increasingly divergent as embryonic development progresses. Additionally, we observed gross disruption of mesoderm diversification beyond purely cardiogenic cell types in Middle- and Late-stage *Mesp1* KO embryos (Fig. 3F-H,K-M, Fig. S8), consistent with their altered morphology.

### Mesp1 knockout cardiac mesoderm cells progress incompletely and imperfectly towards cardiomyocyte fates

We next investigated the steps of cardiac fate progression to understand how *Mesp1* KO embryos initiate cardiogenesis but fail to produce matures CMs. Utilizing pseudotemporal trajectory ordering with URD (Farrell et al., 2018) on the full mesoderm dataset, we defined the epiblast cells, the cluster also containing the earliest staged embryos (C1-Epiblast in Fig. S6E,G), as the root, and clusters containing the most differentiated mesodermal cells from the oldest stage embryos as the tips (Fig. 4A, Fig. S6E,G, Fig. S9A,B). We layered expression of *Smarcd3*-F6-eGFP to identify the main cardiogenic fate paths within the tree space, which also co-expressed CM genes such as *Nkx2-5*, *Myl7*, and *Smarcd3* (Fig. 4C, Fig. S9C). Within CM and CardiacMeso fate branches, *Mesp1* KO cells occupied the youngest pseudotemporal positions near the top of the branch segment, and were more represented in younger pseudotime segment branches of the tree, including their own earlier-pseudotime branch fate “C22” which was defined by multiple mesodermal genes not representative of any particular wildtype cell type (Fig. 4A-B, Fig. S6E,I).

**Fig. 4.**
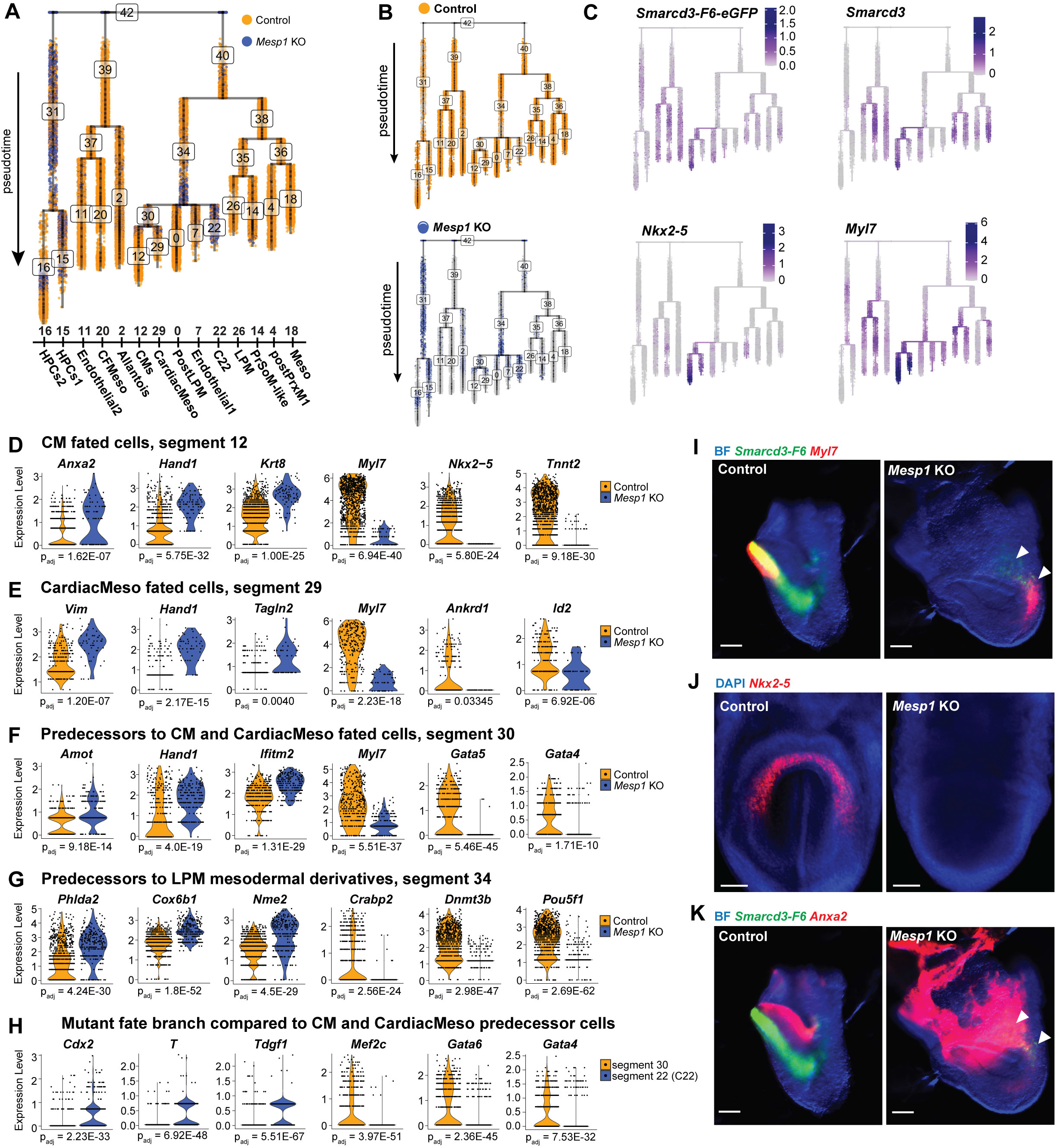
Pseudotime trajectory analysis of mesoderm fates in *Mesp1* KO embryos. (A) URD pseudotime tree for fate progression towards mature mesoderm fates colored by genotypes together and (B) separately. (C) Overlay of cardiac marker gene expression. (D-H) Differentially expressed genes in cells of shared fates and pseudotime identities; (D) CM fated cells, (E) CardiacMeso fated cells, (F) predecessors to CM and CardiacMeso fates, (G) predecessors to LPM derivate fates, (H) comparison of mutant fate branch C22 to predecessors to CardiacMeso fates. (I-J) Multiplexed fluorescence *in situ* hybridization for *Smarcd3*-F6 (green) and (I) *Myl7* (red) and (J) *Nkx2-5* (red), and (K) *Anxa2* in cardiac crescent stage embryos. Arrowheads denote *Smarcd3*-F6+ cardiogenic regions in *Mesp1* KO embryos. Scale bars are 100 μm.

Focusing on the cardiogenic fate tree section beginning at segment 34, we performed differential gene expression analysis to compare cell types of similar fate potentials within branch segments or pseudotemporal levels of the trajectory (Fig. 4D-H). Among CM-fated cells, *Mesp1* KO cells were enriched for expression of *Anxa2*, *Hand1*, *Krt8* and other genes reminiscent of extraembryonic mesoderm, expressed lower levels of structural myocyte genes such as *Myl7* and *Tnnt2* relative to control, and lacked *Nkx2-5* transcripts. (Fig. 4D, Table S5A). *Hand1* was similarly enriched in *Mesp1* KO CardiacMeso-fated cells, along with *Vim*, a fibroblast gene, and *Tagln2*, a gene involved in cell transformation and cell morphology (Han et al., 2017) (Fig. 4E, Table S5B). In the CardiacMeso-fated segment, *Myl7* and *Id2* were reduced relative to controls, as was *Ankrd1*, a gene implicated in sarcomere-binding and dilated cardiomyopathy that is known to be upregulated with overexpression of *Mesp1* (Bondue & Blanpain, 2010; Moulik et al., 2009) (Fig. 4E, Table S5B). In the branch that gave rise to CM and CardiacMeso fates, segment 30, *Amot*, *Hand1*, and *Ifitm2*, genes expressed in the posterior proximal extraembryonic border of the murine embryo and ExEMeso, were increased in *Mesp1* KO cells (Fig. 4F, Table S5C). By contrast, myocyte and cardiac progenitor genes *Myl7*, *Gata5*, and *Gata4* were decreased in *Mesp1* KO cells relative to control (Fig. 4F, Table S5C). In *Mesp1* KO cells in segment 34, the predecessors to LPM mesodermal derivatives, pronephros gene *Cox6b1*, spongiotrophoblast and extraembryonic energy storage gene *Phlda2*, and ESC self-renewal gene *Nme2* (Zhu et al., 2009) were enriched, while retinoic acid gene *Crabp2* and early gastrulation genes *Dnmt3b*, *Pou5f1* were downregulated (Fig. 4G, Table S5D). Finally, we compared the *Mesp1* KO cell-dominated segment 22 to its pseudotime-branching contemporary segment 30, and found enriched expression of mesoderm-fate promoting gastrulation TFs *Cdx2* and *T*, along with mesendoderm allocation gene *Tdgf1* (Fig. 4H, Table S5E). Conversely, cardiac progenitor morphogenesis TFs *Mef2c*, *Gata6*, and *Gata4* were downregulated (Fig. 4H, Table S5E).

We summarize these analyses of *Mesp1* KO cardiac mesoderm fates into two categories; 1) retained expression of some cardiac progenitor genes (*Myl7, Gata4/5/6, Id2, Tnnt2*), albeit at decreased levels relative to control, and absence of others (*Nkx2-5, Ankrd1)*, and 2) ectopic enrichment of ExEMeso and other mesoderm associated genes (*Hand1, Anxa2, Amot, Vim, Tagln2*). We used multiplexed fluorescent RNA *in situ* hybridization to validate the spatial domains of differentially expressed genes in Late-stages, and confirmed presence of *Myl7*+ cells co-expressing *Smarcd3*-F6 in the posterior distal compartment of *Mesp1* KO embryos (Fig. 4I) along with absence of *Nkx2-5* expression in *Mesp1* KO embryos (Fig. 4J). We also showed ectopic *Anxa2* expression into the embryo proper, overlapping with *Smarcd3*-F6+ cells in their posterior position in *Mesp1* KO embryos, in contrast to the anterior extraembryonic-restricted expression pattern of controls (Fig. 4K). These results further highlight that *Mesp1* KO CPCs ectopically express non-cardiac mesodermal genes, and reveals that *Mesp1* KO CPCs progress towards CM fates incompletely in part through a failure to express requisite TFs. Thus, *Mesp1* KO CPCs reach a cardiogenic breakpoint during gastrulation prior to cardiac crescent formation.

### scATAC-seq analysis reveals a regulatory barrier in *Mesp1* KO mesoderm progression towards cardiomyocyte fates

To characterize the regulatory landscape prohibiting *Mesp1* KO cells from progressing fully towards CM fates, we turned to single cell Assay for Transposase Accessible Chromatin (scATAC-seq) (Buenrostro et al., 2015) of Middle- and Late-stage embryos ages E7.5 - E7.75 (Fig. S10). We processed whole embryos and performed preliminary atlasing analysis in ArchR (Granja et al., 2021). We utilized integration with the complementary whole embryo scRNA-seq dataset along with chromatin accessibility profiles near marker genes (gene scores) to subset mesodermal cell type clusters (Fig. S11A-D, Table S6A) in order to generate a subset scATAC-seq atlas of 16 mesodermal cell types (Fig. 5A). *Mesp1* KO and controls had strikingly divergent regulatory landscapes (Fig. 5B). *Mesp1* KO cells were confined to scATAC-seq clusters representing epiblast (Epi), mesendoderm, and LPM cell types, while control cells were represented in the LPM cell types, the more mature cardiac progenitor (CP) and CM cluster, and mesodermal derivative cell types (Fig. 5A-C). Integration with the complementary mesoderm scRNA-seq dataset (Table S6B), visualization of key marker gene scores and integrated expression (Fig. 5D-E), and Jaccard indexing (Fig. S12) were used to assign relative cell identities to each mesoderm scATAC-seq cluster (Fig. 5C). While some cardiac TFs such as *Nkx2-5* were not active in *Mesp1* KO cells, others such as *Tbx5* had chromatin accessibility in *Mesp1* KO cells, but integrated expression only in control CM/CP cells (Fig. 5B,D-E). Other cardiac TFs *Hand1* and *Gata4* had similar activity between *Mesp1* KO and control cells (Fig. 5B,D-E), and while *Mesp1* KO cells downregulated *Smarcd3* and *Myl7* expression, chromatin accessibility for these genes was similar between genotypes (Fig. 5B,D-E). These results indicate a perdurance of active chromatin states in the steps preceding cardiogenic differentiation.

**Fig. 5.**
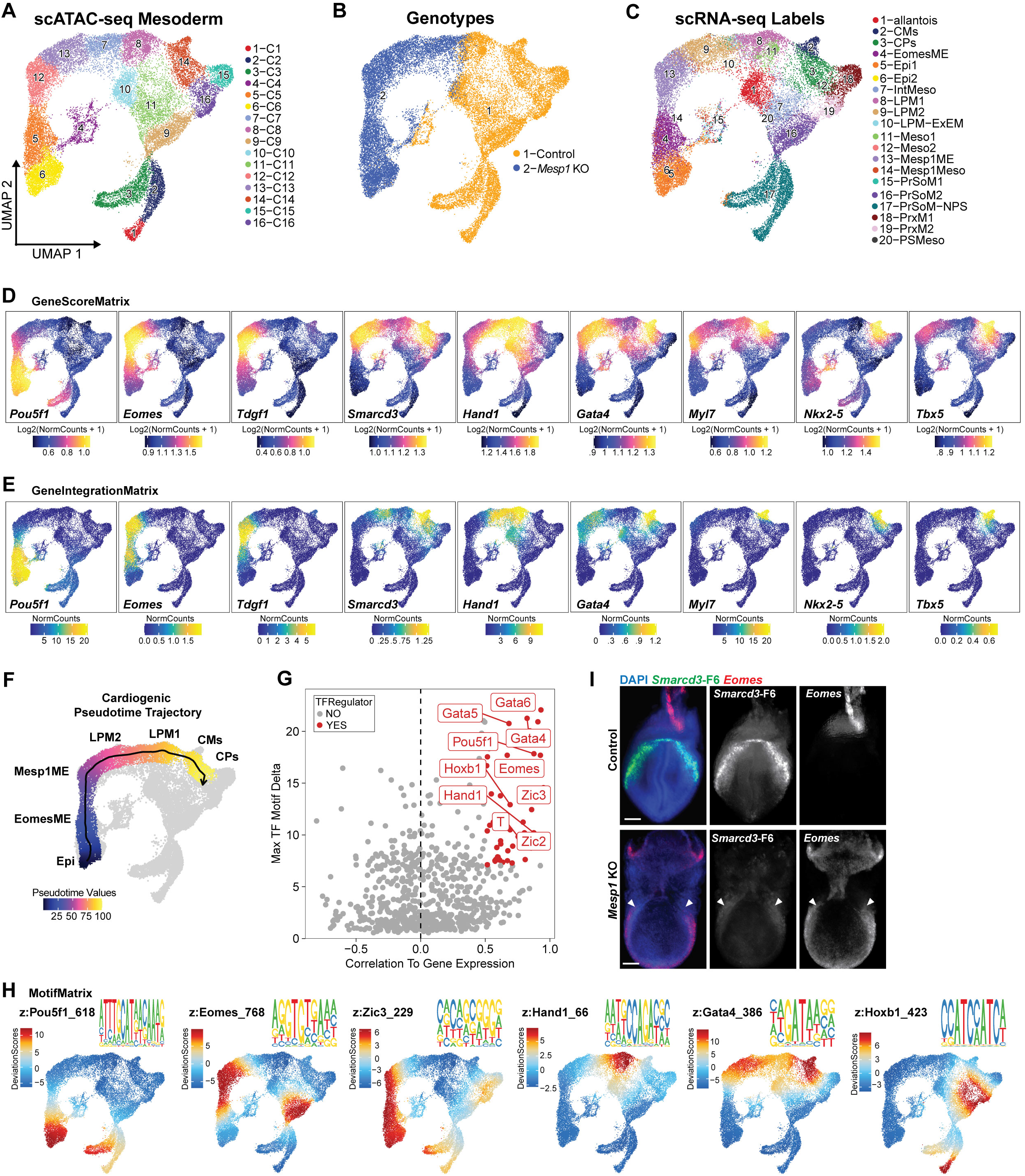
Characterizing transcriptional drivers in *Mesp1* KO mesoderm during cardiogenesis. (A) Mesoderm scATAC-seq atlas of 16 cell types with overlays for (B) genotypes and (C) relative cell type identities from integration of a complementary scRNA-seq dataset. (D) GeneScoreMatrix plots for chromatin accessibility around gene loci and (E) GeneIntegrationMatrix plots for scRNA-seq integrated gene expression for cardiac mesoderm marker genes and TFs. (F) Pseudotime values for cells along the *Mesp1* KO cardiac-fate trajectory path. (G) Maximum z-score delta for TF motif variance between clusters correlated to gene expression within clusters to identify positive TF drivers (red). (G-H) Highlighted positive regulator TFs’ motif z-scores mapped in UMAP space, with associated position weight matrix plots. (I) Multiplexed fluorescence *in situ* hybridization for *Smarcd3*-F6 (green) and *Eomes* (red) in cardiac crescent stage embryos. Arrowheads denote cardiogenic regions in *Mesp1* KO embryo. Scale bars are 100 μm.

To interrogate the developmental relationship between *Mesp1* KO cells failing to mature and control CMs, we performed an ArchR trajectory inference analysis assessing pseudotime along the cardiac fate path. We defined a trajectory backbone in the *Mesp1* KO cells traversing the expected differentiation path of Epi, *Eomes*+ mesendoderm (EomesME), *Mesp1*+ mesoendoderm (Mesp1ME), lateral plate mesoderm (LPM2, LPM1), to cardiac progenitors (CP) and cardiomyocytes (CM) clusters. This trajectory analysis revealed that while *Mesp1* KO cells traversed the normal path from epiblast to LPM, they abruptly failed to progress further towards CPs and CMs (Fig. 5F). Notably, the most mature cell identity *Mesp1* KO cells achieved (LPM1) also contained control cells capable of progressing to CPs past this point where *Mesp1* KO cells halted, indicating the LPM1-to-CP transition represents the breakpoint in cardiogenesis for *Mesp1* KO cells (Fig. 5F).

From this trajectory analysis, we assessed dynamic shifts in the correlation of TF gene scores and gene expression with corresponding TF motifs in accessible chromatin peaks across pseudotime (Fig. S13A-B) to reveal a biologically-sensical order of TF regulators involved in cardiogenesis. Notably, TFs represented in early pseudotime and *Mesp1* KO cells (*Lhx1, T, Eomes, Zic2/3, Pitx2, Isl1*, Fig. S13A-B) were consistent with early gastrulation mesodermal regulatory networks, indicating that aspects of these networks are either *Mesp1*-independent or resilient to *Mesp1* loss. TFs represented in later pseudotime (*Hand2, Gata4/5/6, Hoxb1*, Fig. S13A-B) were concordant with downregulated gene expression in *Mesp1* KO CPCs and mesoderm by scRNA-seq (Fig. 2-4), suggesting that failed induction of these TFs and their programs is either *Mesp1*-dependent or vulnerable to secondary effects of *Mesp1* loss.

To ascertain which gene regulatory networks were present in which cell types, and thus which genotypes, along the cardiogenic trajectory, we performed an orthogonal analysis to identify putative positive transcriptional drivers (Fig. 5G, Table S6C) and visualized resulting TFs’ motif enrichments in UMAP space (Fig. 5H, S13A-B). In particular, the *Mesp1* KO Epi cluster is driven in part by pluripotency TFs Pou5f1 and Mesp1-cofactor Zic3 (Lin et al., 2022) (Fig. 5D-E, 5H). Mesendoderm TFs Eomes and Zic3 were drivers of EomesME and Mesp1ME (Fig. 5D-E, 5H). ExEM and first heart field TF Hand1 appeared in the “last-stop” LPM1 cell types where *Mesp1* KO cells failed to progress towards more mature cardiac fates (Fig. 5D-E, 5H), consistent with the upregulated expression observed in *Mesp1* KO CM-fated cells (Fig. 4D-F). While Gata motifs were present in LPM2 and LPM1, the latter of which contains both genotypes, Gata4 was most enriched in the later cardiac fate destinations of CPs and CMs (Fig. 5H). This result coupled with the *Gata4*’s representation in late trajectory pseudotime (Fig. S13A) and downregulated expression in cardiac-fated *Mesp1* KO mesoderm cells (Fig. 4F) likely signifies *Mesp1*-dependent induction and/or influence of Gata factor-associated networks within emerging CMs. Indeed, *Gata4* was shown to be activated by Mesp1 during *in vitro* differentiation (Soibam et al., 2015), and while Gata4 binds the minority of Mesp1-bound enhancers, Gata4 binds nearly half of enhancers opened following *in vitro* induction of Mesp1 (Lin et al., 2022). Separately, Mesp1 target gene *Hoxb1*’s motif was distinctly expressed in PrSoM cell types, coincident with the “late phase” role for *Mesp1* (Lin et al., 2022; Haraguchi et al., 2001) in mesoderm diversification beyond the cardiac lineage (Fig. 5H). The preponderance and accordance of these results supports that early cardiogenic phases proceed resilient to *Mesp1*-loss, however *Mesp1* KO cells cannot proceed to later phases.

Given enrichment of Eomes motifs, gene score, and gene expression in mesendoderm clusters (Fig. 5D-E, 5I), its apparent role as a positive TF driver (Fig. 5G,I) and its direct involvement in Mesp1 induction (Tosic et al., 2019; Costello et al., 2011; Alexanian et al., 2017; Guo et al., 2018; Probst et al., 2020), we investigated Eomes as a potential driver of *Mesp1-*independent early phases of cardiogenesis. Eomes directly binds *Myl7* regulatory regions (Tosic et al., 2019), and *Eomes* loss disrupts induction of *Myl7* (Costello et al., 2011), supporting that expression of *Myl7* in *Mesp1* KO CPCs (Fig. 2E,K, Fig. S5A,C-E, Fig. 3D, Fig. 4C-F,J) is regulated by Eomes at least partially independently of Mesp1. Furthermore, domains of *Eomes* expression anomalously endured in cardiogenic *Smarcd3*-F6+ regions and are ectopically expanded in lateral aspects of the embryo proper in cardiac-crescent staged *Mesp1* KOs (Fig. 5I), indicating improper repression of *Eomes* in cardiogenic regions.

Taken together, these results describe a shift between mesendoderm and cardiac patterning regulatory programs during cardiogenesis. *Mesp1* KO cells are unable to traverse beyond LPM cell types to initiate cardiac patterning programs and instead retain gene expression indicative of earlier cardiac mesoderm regulatory programs. The perdurant expression of *Eomes* may be a driving mechanism for this halt in cardiogenesis.

### The disrupted regulatory landscape of *Mesp1* KO embryos is characterized by ectopic endurance of mesendoderm gene programs

To understand how the *Mesp1* KO disrupted regulatory landscape underlies the transcriptional barriers to progression towards more mature cardiac fates, we characterized cell type peak accessibility profiles and the motif enrichment within these peaks (Fig. S14A-B). We performed differential accessibility testing of peaks between cell types along the cardiac trajectory (Fig. S14C-G). Focusing specifically on the “last stop” for *Mesp1* KO cells, we compared motif enrichment within differential peaks of CMs/CPs containing only control and LPM1 containing both control and *Mesp1* KO cells (Fig. 6A). In agreement with motif enrichment scores for positive TF regulators (Fig. 5G, 5H), Gata and Mef2c motifs were among those enriched in the more mature cardiac fates, while motifs for cardiac differentiation and myogenesis-promoting Tead factors were relatively enriched within LPM1, offering further explanation for retention of some myocyte identity within *Mesp1* KO CPCs (Fig. 6A, Fig. 2E, 2K, Fig. S5D, Fig. 4J) (Han et al., 2020; Akerberg et al., 2019). To measure correlations between the differential accessibility profiles behind these motifs and the complementary gene expression profiles of these cells, we performed an association analysis measuring the probability that peak accessibility near genes corresponds to gene expression. We applied this analysis to find an odds ratio of 16.7 for the probability that significantly differentially open peaks corresponded to upregulated gene expression (Q3, Fig. 6B) while significantly differentially closed peaks corresponded to downregulated gene expression (Q1, Fig. 6B) in CMs/CPs relative to LPM1. Thus, gene expression profiles enriched in control-only CMs/CPs (Q3: *Mef2c, Tbx5, Gata5, Nkx2-5, Tnnt2*, Fig. 6B) and transcriptional profiles of LPM1 cells (Q1: *Hand1*, *Anxa2, Cdx2, Krt8/18*, Fig. 6B) are associated with these cells’ differing chromatin landscapes.

**Fig. 6.**
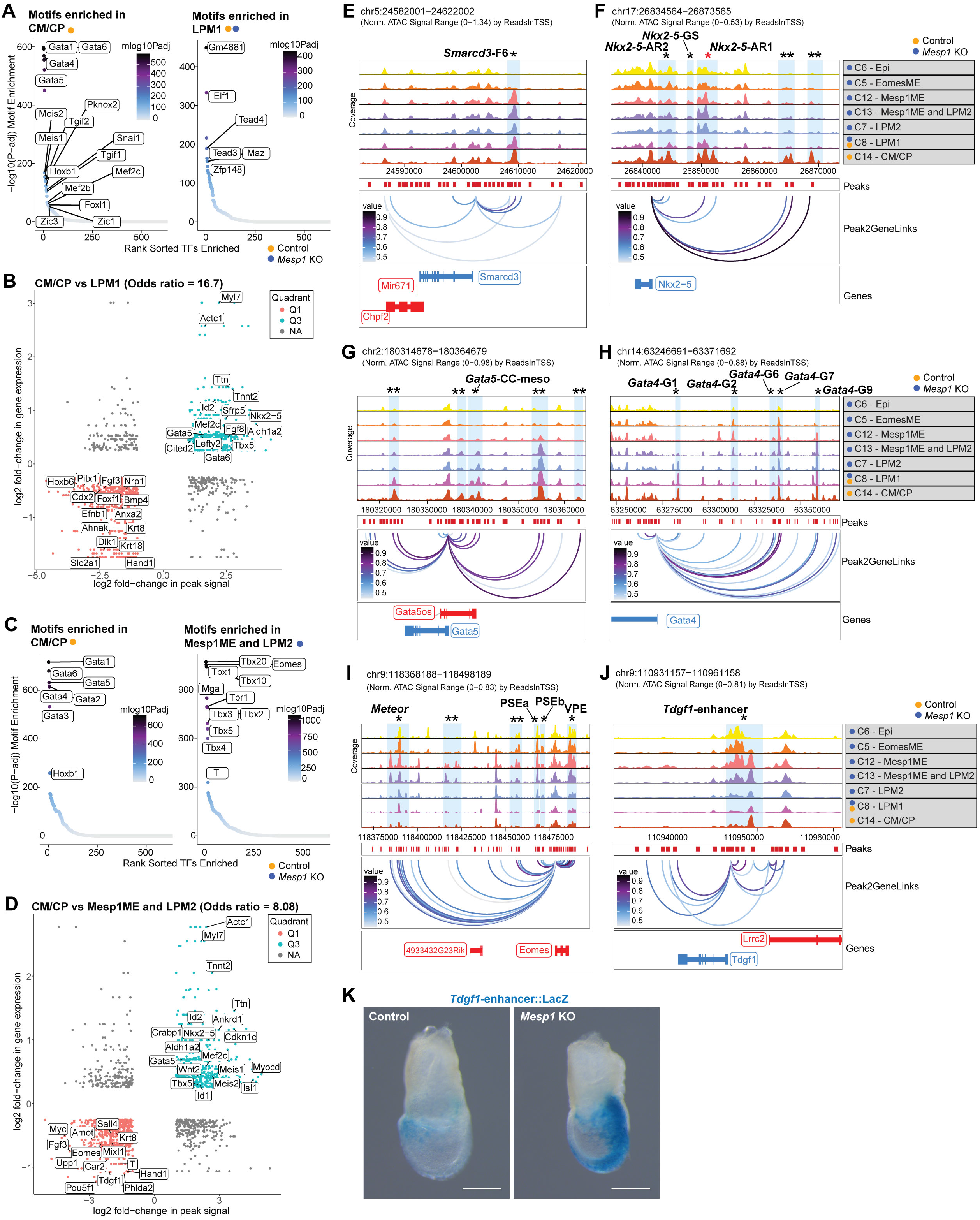
Disrupted regulatory landscape of *Mesp1* KO mesoderm. (A,C) Motifs enriched in differentially accessible peaks between (A) CM/CP vs. LPM1 and (C) CM/CP vs Mesp1ME and LPM2. (B,D) Plots for peak,gene associations showing correlations between differential peak accessibility and gene expression in comparisons between cells type1 vs type2. Q3 peak,gene pairs represent significantly more accessible peaks paired with upregulated gene expression in type1 cells. Q1 peak,gene pairs represent significantly more accessible peaks paired with upregulated gene expression in type2 cells. Odds ratio denotes probability for observed peak,gene relationships. (B) Peak,gene association plot for cells in CM/CP vs LPM1 comparison and (D) in CM/CP vs Mesp1ME and LPM2 comparison. (E-J) Peak2Gene linkage browser tracks for cell types showing predicted regulatory connections between distal accessible regions (Peaks) and nearby genes. Shaded bars denote predicted distal regulatory regions; *denotes characterized elements; red* denotes regions with Mesp1-binding; **denotes uncharacterized elements. Characterized elements named when available. (E) *Smarcd3* linkage to the “F6” enhancer. Peak linkages to genes (F) *Nkx2-5*, (G) *Gata5*, (H) *Gata4*, (I) *Eomes*, and (J) *Tdgf1.* (K) X-gal stain for activity of characterized *Tdgf1* enhancer, scale bars are 200 μm.

We next compared control-only CMs/CPs to *Mesp1* KO-only Mesp1ME and LPM2 (Fig. 6C-D) because these cells had similar gene scores for the *Smarcd3* locus (Fig. 5D), a proxy for *Smarcd3*-F6 enhancer activity. Motifs including those for Gata and Hox factors were relatively enriched in CMs/CPs, and T-box motifs including Eomes and T were enriched in Mesp1ME and LPM2 (Fig. 6C). The correlation odds ratio of 8.08 highlighted corresponding peak accessibility and gene expression enrichment for CP patterning and CM genes in control CMs/CPs (Q3: *Nkx2-5, Tbx5, Wnt2, Mef2c, Meis1, Ttn, Tnnt2*) and relative enriched peak accessibility near upregulated genes for earlier cardiac mesoderm and mesendoderm programs in Mesp1ME and LPM2 *Mesp1* KO-only cells (Q1: *Tdgf1, Fgf3, Eomes, Mixl1, T, Krt8, Hand1, Pou5f1*) (Fig. 6D).

Applying this analysis paradigm to multiple pairwise comparisons along the cardiogenic trajectory (Fig. S14H-M) showed that the predominant regulatory signature of control CMs/CPs, is characterized by TFs such as *Gata4/5/6, Hoxb1, Mef2c, Foxf1, and Tbx5*, which are required for initiation of cardiac patterning and morphogenesis programs upon formation of the cardiac crescent, subsequent heart fields, and higher level organogenesis (Pikkarainen et al., 2004; Kokkinopoulos et al., 2015; Bruneau, 2013; Stefanovic et al., 2020; Harvey, 2002; Kelly, Buckingham & Moorman, 2014). *Mesp1* KO cells were unable to activate these same regulatory programs, instead retaining TFs for mesendoderm and other mesoderm networks (*T, Eomes, Hand1)* (Fig. S14H-M).

To visualize regulatory interactions between chromatin accessibility and integrated gene expression agnostic of differential accessibility and expression testing between specific cell types, we utilized the ArchR pipeline’s orthogonal “peak2gene” linkage approach (Granja et al., 2021). This linkage prediction method identified both known and uncharacterized distal regulatory elements (Fig. 6E-J). The *Smarcd3*-F6 enhancer (Devine et al., 2014) expectedly showed linkage to *Smarcd3* and similar accessibility across cardiogenesis, including the *Mesp1* KO cells Mesp1ME, LPM2 (Fig. 6E), consistent with our detection of the transgene by scRNA-seq.

Accordant with the modular enhancer landscape of *Nkx2-5*, multiple peak linkages were defined for the *Nkx2-5* locus, including two distal uncharacterized regions (Fig. 6F). Two linkages were appropriately mapped to the characterized Gata4-, Nfat-, Mesp1/Mzf1-, and Isl1-regulated 9 kb-upstream *Nkx2-5* cardiac enhancer sequence (*Nkx2-5*-AR1) (Lien et al., 1999; Chen & Cao, 2009; Clark et al., 2013; Doppler et al., 2014; Bondue et al., 2008) and the distal-linked AR1 peak was increased in control CM/CP cells only (Fig. 6F). Similarly, the Gata-, Smad4-, Nfat-, Isl1-regulated *Nkx2-5*-AR2 enhancer (Searcy et al., 1998; Liberatore et al., 2002; Lien et al., 2002) and the two uncharacterized linked regions ∼25 kb and ∼30 kb-upstream of the TSS showed enriched accessibility in control CM/CP cells (Fig. 6F) while Gata4- and Smad1/4-responsive 6 kb-upstream enhancer (*Nkx2-5*-GS) (Brown et al., 2004) didn’t show accessibility in any cells (Fig. 6F). These results are consistent with absence of *Nkx2-5* in *Mesp1* KO embryos (Fig. 4), underscore the complexity of regulation on this critical cardiac TF, and provide further evidence for the regulatory shift between LPM1 and CM/CP cells (Fig. 5F) that *Mesp1* KO cells are unable to progress through.

Examination of the *Gata5* locus revealed a linkage to the characterized cardiac crescent and mesodermal derivatives enhancer (*Gata5*-CC-meso) (MacNeill et al., 2000) as well as several uncharacterized linked distal elements with accessibility in Mesp1ME, LPM2, LPM1, and CM/CP cells (Fig. 6G). Several characterized *Gata4* enhancer regions were linked, including lateral mesoderm enhancer *Gata4*-G2 (Rojas et al., 2005) and cardiac crescent enhancer *Gata4*-G9 (Schachterle et al., 2012). Foxf1 and Gata4-bound enhancer *Gata4*-C2 showed enriched accessibility in mesendoderm *Mesp1* KO cells, while ETS-activated *Gata4*-G9 was similarly accessible between between Mesp1ME, LPM2, LPM1, and CM/CPs (Fig. 6H), highlighting retention of active chromatin states preceding cardiac patterning and differentiation despite loss of *Mesp1*.

Evaluation of loci for mesendoderm genes *Eomes* and *Tdgf1,* which ectopically perdure in *Mesp1* KO embryos, showed a corresponding pattern of enriched linked peaks in *Mesp1* KO cell types (Fig. 6I). The characterized distal element *Meteor*, a lncRNA (Alexanian et al., 2017), was linked to *Eomes* with greatest accessibility enrichment in Epi, EomesME, and Mesp1ME *Mesp1* KO cells (Fig. 6I). Similarly, characterized PSEa, PSEb, and VME regulatory regions (Simon et al., 2017) were linked (Fig. 6I), supporting that the early cardiac mesoderm transcriptional landscape is intact despite *Mesp1* absence, however retained later in development than it should be for the age of these embryos. Upstream of *Tdgf1*, a previously characterized enhancer sequence and direct transcriptional target of Mef2c (Barnes et al., 2016) displayed enrichment of proximal peaks in *Mesp1* KO cells (Fig. 6J), which we confirmed by increased *Tdgf1* enhancer transgene activity in posterior domains of E7.5 *Mesp1* KO embryos (Fig. 6K). Increased *Tdgf1* enhancer activity mimicked the enriched *Tdgf1* gene expression in *Mesp1* KO embryos (Fig. 2F-G), further supporting the hypothesis that early programs are de-repressed in absence of *Mesp1*.

We examined additional linked peak profiles around differentially expressed genes (Fig. S15). We detected linkages between *Mesp1* and the characterized “EME” enhancer (Haraguchi et al., 2001; Ajima et al., 2021; Costello et al., 2011; Guo et al., 2018) with enrichment in *Mesp1* KO cell types likely indicative of retained early chromatin landscape or de-repression of the locus without appropriate regulation from downstream targets (Fig. S15A). We detected linkages to 3 uncharacterized distal regions near the *Gata6* locus, as well as the Nkx2-5-targeted enhancer regions (Molkentin et al., 2000) which had similar accessibility profiles across *Mesp1* KO Mesp1ME, LPM2 cells, and the LPM1 cells containing both genotypes (Fig. S15B). We noted linkages to multiple characterized *Hand1* enhancer regions (Vincentz et al., 2021, 2019; George & Firulli, 2021) across both genotypes and multiple cell types, with accessibility for some enhancers decreasing in CM/CP cells (Fig. S15C), consistent *Hand1*’s more robust activity in LPM1 cells (Fig. 5D,E,H). Peaks with similar accessibility across Mesp1ME and LPM cell types containing both genotypes were detected in linkages near *Tbx5,* including the Tbx5-CRE16 (Smemo et al., 2012), and downregulated but retained structural myocyte genes *Tnnt2* (Fig. S15E) and *Myl7* (Fig. S15F). Increased accessibility for *Anxa2-*linked peaks in *Mesp1* KO LPM2 cells and control/*Mesp1* KO LPM1 cells contrasted near-inaccessibility in control CM/CPs (Fig. S15G), consistent with the upregulated *Anxa2* expression in Late-stage embryo cardiogenic regions (Fig. 4K). Downstream distal peaks were linked to Mesp1-induced EMT-gene *Snai1* (Fig. S15H) (Lin et al., 2022), including the Mesp1-binding site. These peak2gene linkage analyses further illustrate the correlation between differentially expressed genes and the altered chromatin landscape in *Mesp1* KO mesoderm cells that prevents progression towards mature cardiac fates, but also highlights that despite this disrupted regulatory landscape, some distal-regulatory elements relevant to early cardiogenesis are still retained.

## DISCUSSION

We generated scRNA-seq and scATAC-seq datasets from whole mouse embryos in a timeline of gastrulation, creating a valuable *in vivo* resource for high-resolution studies of gene regulatory networks in early embryonic development. We utilized computational detection of the CPC-labeling transgenes to focus on early cardiac specification, showing that while *Mesp1* KO embryos are capable of initiating and progressing through early cardiac mesoderm specification, a *Mesp1*-dependent regulatory barrier prevents *Mesp1* KO CPCs from progressing completely towards CM fates. We characterized improper repression of early mesendoderm programs at this breakpoint, such as how absence of *Mesp1* leads to enduring *Eomes* activity, which in turn promotes ectopic perdurance of mesendoderm transcriptional networks when cardiac crescent-staged embryos should instead be upregulating cardiac patterning programs. Additionally, this disrupted regulatory landscape likely contributes to *Mesp1* KO cardiac mesoderm and CPCs ectopically expressing non-cardiac mesoderm genes. Despite this ectopic expression, CPCs do not appear to deviate from a cardiac-directed mesodermal fate path. Ultimately, while *Mesp1* KO embryos specify early cardiac lineage cell types, their deficient regulatory landscapes prove prohibitive against further lineage development (Fig. 7).

**Fig. 7.**
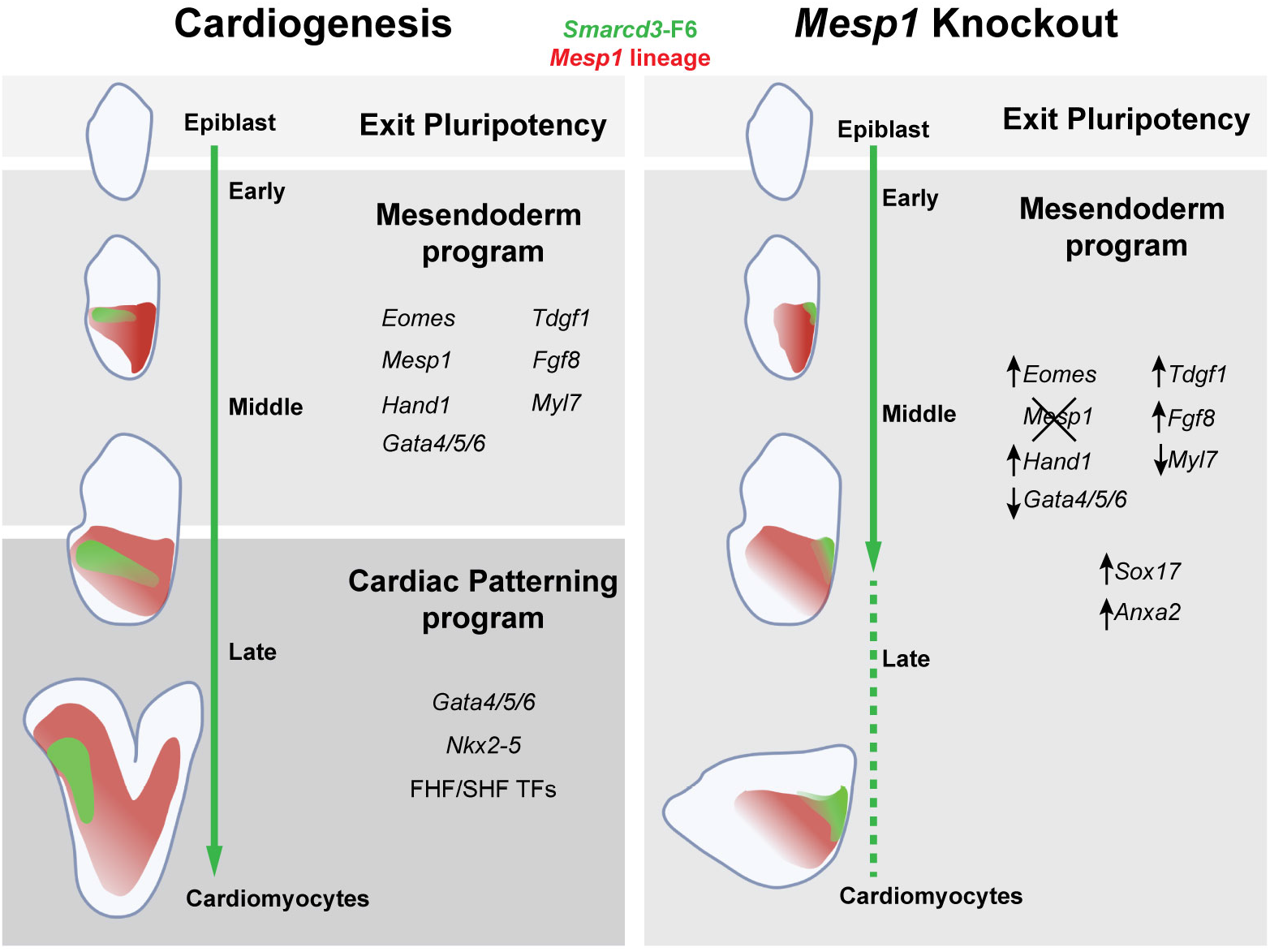
Model for transcriptional regulatory landscape of cardiogenesis and loss of *Mesp1*. Schematic model of gene regulatory program phases during cardiac mesoderm specification and differentiation. *Mesp1* KO cardiac mesoderm cells exit pluripotency, induce early cardiac specification genes under control of mesendoderm programs, yet fail to activate critical cardiac TFs at cardiac crescent stages to initiate cardiac patterning programs.

Positing *Mesp1* as a master transcriptional regulator of early cardiac fate is largely informed by overexpression studies (Chiapparo et al., 2016; Bondue et al., 2008; Lindsley et al., 2008; Wu, 2008; Kelly, 2016; Bondue & Blanpain, 2010; Lin et al., 2022) in contrast to earlier *in vivo* studies which suggested a *Mesp1*-dependent role for cardiac mesoderm migration (Saga, Kitajima & Miyagawa-Tomita, 2000). Indeed, in this work we note downregulation of migratory genes in *Mesp1* KO cells, and a companion work demonstrates that *Mesp1*-dependent migration patterns are critical for spatial organization of CPCs during cardiogenesis (Dominguez et al., 2022). Additional interpretations in *Mesp1/Mesp2* double knockouts underscore the potential for more complex networks of TF dependency in cardiac specification not fully explained by regulatory hierarchies (Ajima et al., 2021; Kitajima et al., 2000; Saga, 1998). While the concept of a “master transcription factor” is a broadly applied hierarchical framework for interrogation of gene regulatory networks (Cai et al., 2020; Davis & Rebay, 2017; Yin & Wang, 2014), and *Mesp1*’s coincident expression in emerging CPCs supports an instructive role for *Mesp1* in cardiogenesis, this model likely oversimplifies cardiogenesis. Indeed, our high resolution, single cell transcriptional and epigenomic analyses reveal both transcriptional resilience and vulnerability of early cardiogenesis in a regulatory landscape lacking *Mesp1*.

Transcriptional profiling of *Smarcd3*-F6+ cells highlighted that *Mesp1* KO cells were mostly represented in cell types of early cardiogenesis and in Early- and Middle-stage embryos, indicating that *Mesp1* KO CPCs not only initiate but also progress through early stages of cardiac specification. This finding contrasts with the *Mesp1*-dependent failure to exit pluripotency previously highlighted (Lescroart et al., 2018). We interpret the failed induction of *Nkx2-5*, which is critical for patterning of the first and second heart fields (Harvey, 2002), and the inappropriate levels of *Gata* factors in *Mesp1* KO CPCs as representing a breakpoint between phases of the cardiogenic process.

To characterize this breakpoint, we utilized complementary scATAC-seq and scRNA-seq mesoderm datasets to conclude that mesendoderm regulatory programs, instructed at least partially by *Eomes,* are responsible for the initiation and progression through Middle phases of cardiac specification prior to cardiac crescent formation. However, the perdurance of these programs coupled with the failure of LPM to properly migrate anterior-laterally in *Mesp1* KO embryos leads to aberrant upregulation and ectopic expression of early cardiac mesoderm, non-cardiac mesoderm, and mesendoderm genes and TFs. Additionally, we hypothesize that the posterior positioning of CPCs in *Mesp1* KO embryos further compounds cardiac maturation and CPC transcriptional profiles via improper exposure to signaling gradients and growth factors. The dysregulated identity of *Mesp1* KO cardiac mesoderm in this phase between Middle and Late embryonic stages stalls *Mesp1*-deficient cardiogenesis due to failed induction of cardiac progenitor patterning, morphogenesis, and CM maturation regulatory programs.

Although this developmental breakpoint is observed between E7.5-E7.75, well after transient *Mesp1* expression has declined, these processes appear to be *Mesp1*-dependent. Possible explanations for this phenomenon are 1) improper repression of earlier regulators, such as *Eomes*; 2) compounded, *Mesp1*-dependent secondary effects influencing de-repression or ectopic activation; or 3) *Mesp1* KO CPCs are exposed to improper embryonic signaling cues as a result of their aberrant posterior localization. Future studies with additional genetic models and assays of embryos representing earlier developmental timepoints are needed to disentangle these possibilities.

Overall, our work shows that complex transcriptional networks and interdependent hierarchies govern CPC emergence and differentiation. We characterize an initial, transcriptionally resilient, phase of CPC specification and identify that the epigenomic landscape necessary for CPCs to transition from LPM to CPs and CMs is dependent on upstream Mesp1 activity. Our results point to generalizable transcriptional regulatory principles during gastrulation for the allocation of precursor cells from embryonic germ layers towards restricted fates, and differentiation to distinct functional cell types.

## SUPPLEMENTAL FIGURE LEGENDS

**Fig. S1.**
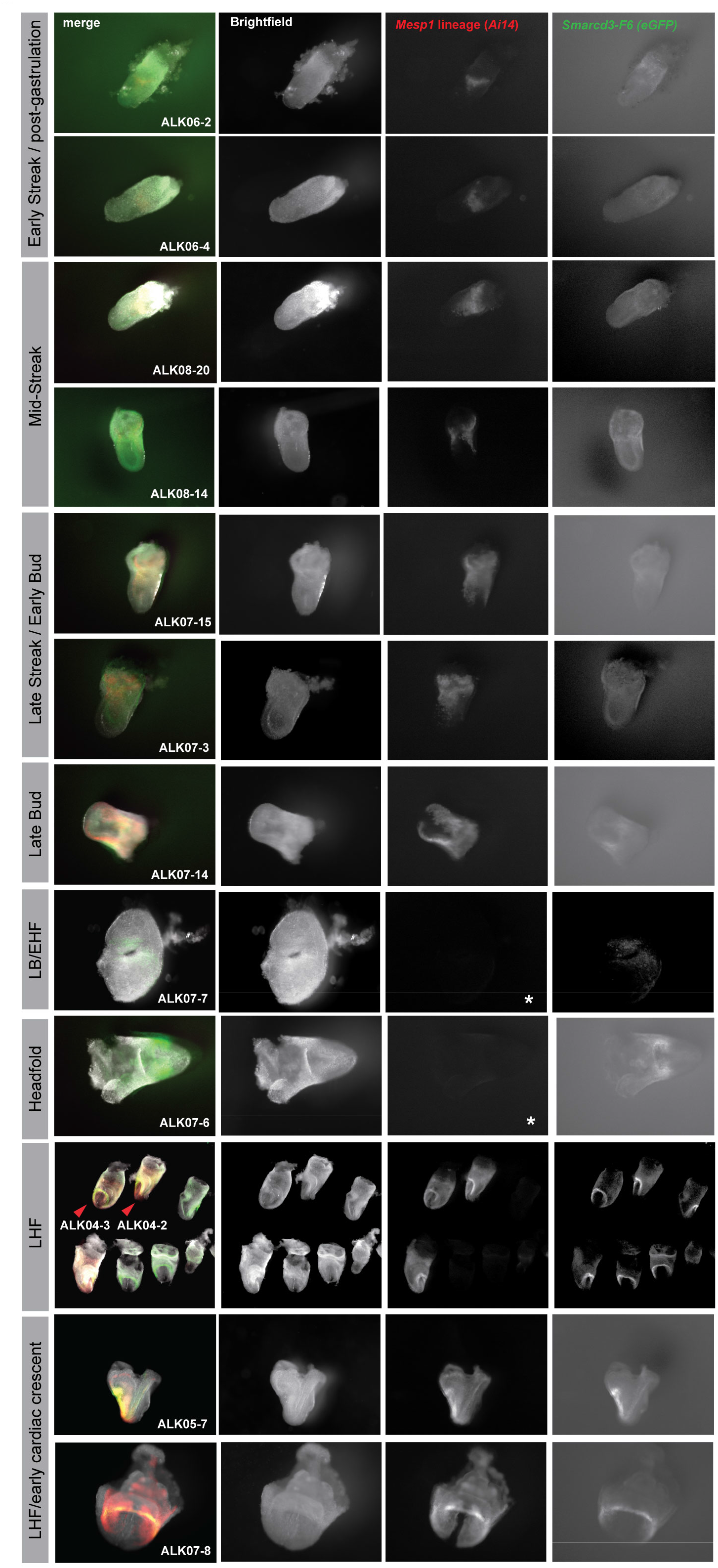
Fluorescent lineage transgenes in whole embryos. Images of all embryos utilized in generation of wildtype gastrulation atlas. *Mesp1* lineage visualized by Ai14 fluorescent reporter transgene. *Smarcd3*-F6 visualized by eGFP fluorescent reporter transgene. Images not acquired and processed identically. Embryos distingushed with * lacked *Mesp1* lineage tracing by Ai14 transgene.

**Fig. S2.**
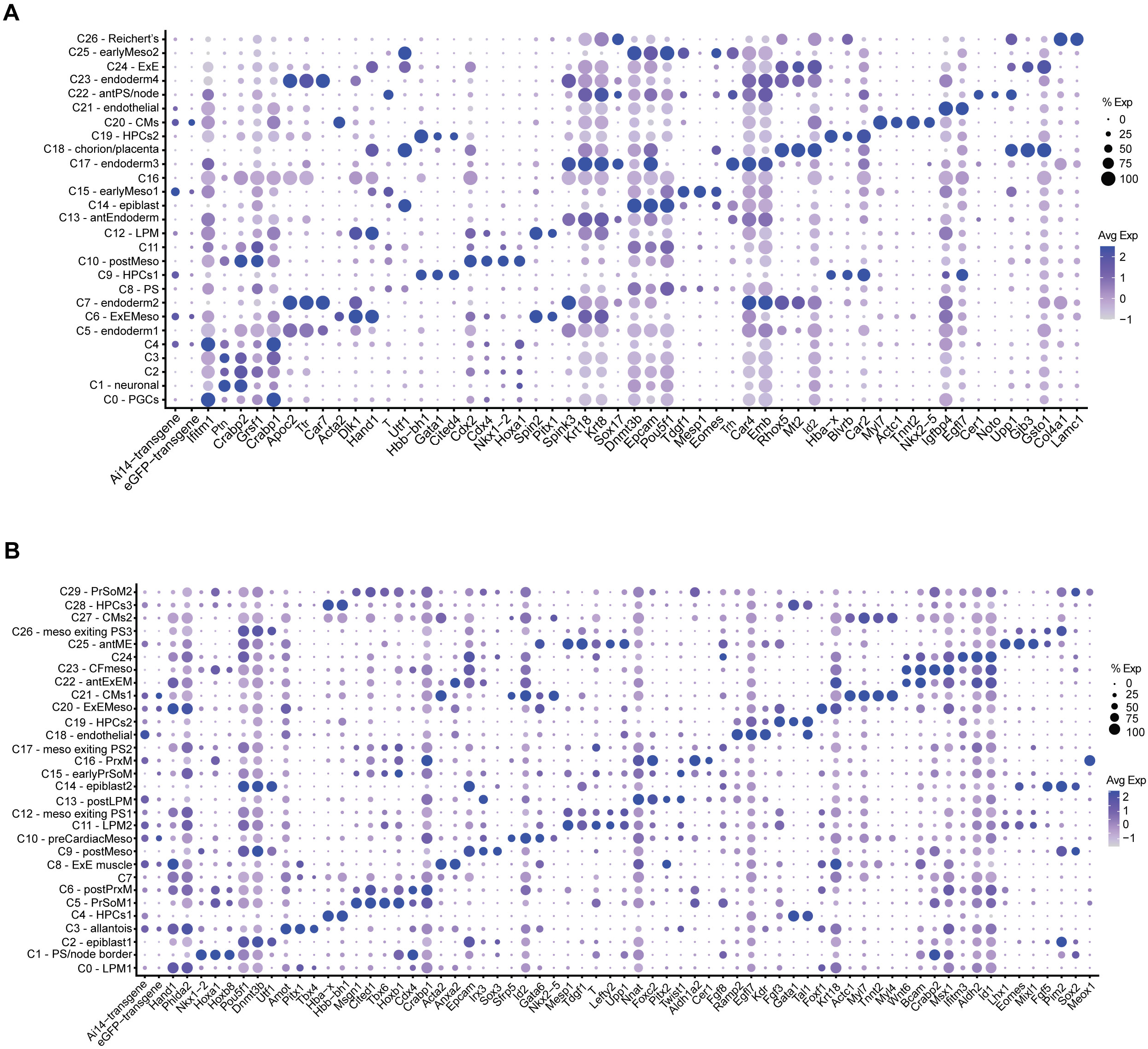
Cell type cluster gene expression profiles. (A) Dotplot denoting marker genes and cell type annotations by cluster in full embryo wildtype gastrulation atlas. (B) Doplot denoting marker genes and cell type annotations by cluster in mesoderm wildtype atlas. Size of dot represents percent of cells expressing gene and color represents average expression level. Cluster number used to denote cell types when annotation was not possible.

**Fig. S3.**
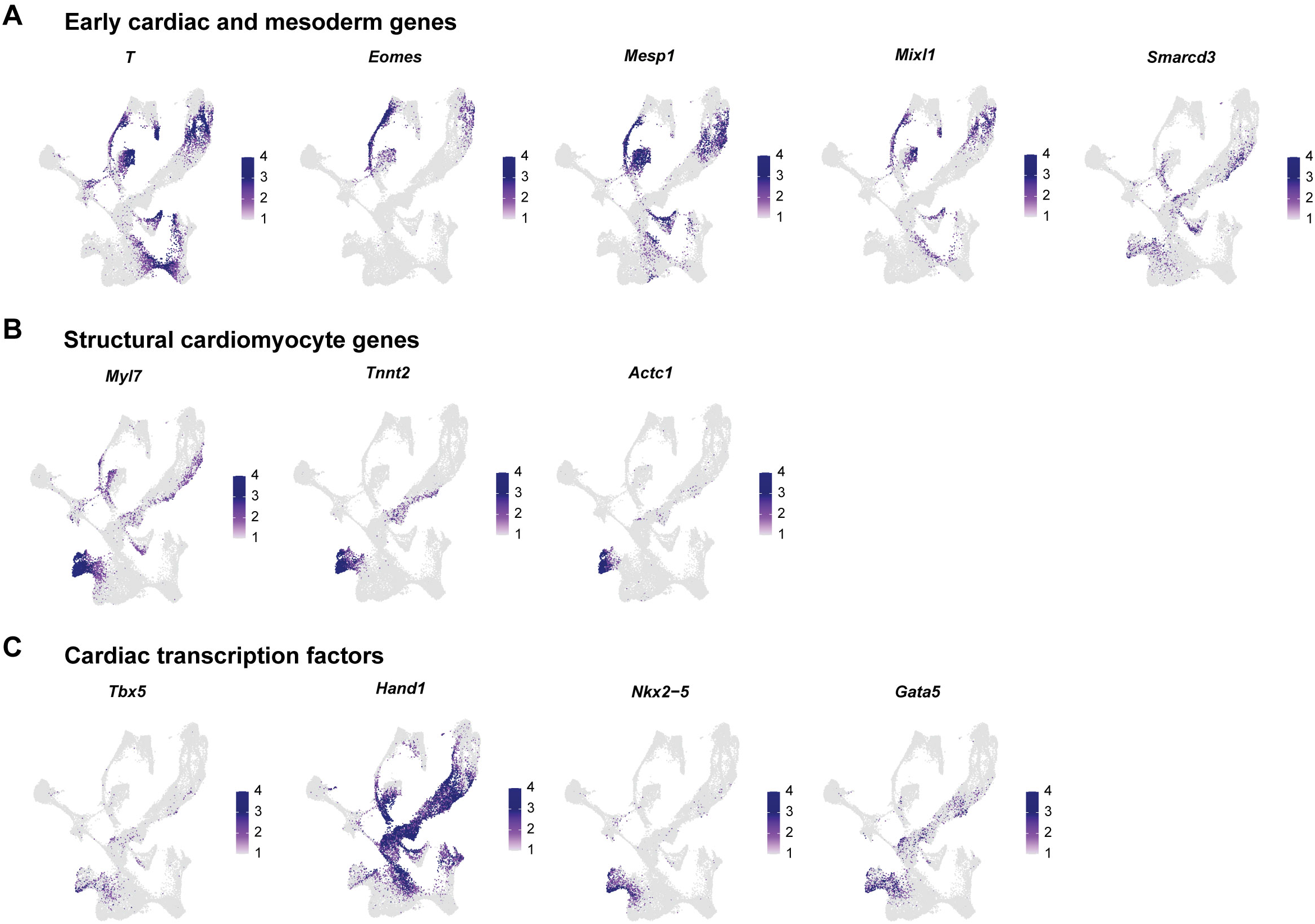
Co-expression of cardiac mesoderm genes in *Smarcd3*-F6+ cell types. UMAP feature plots of wildtype mesoderm atlas showing gene expression of (A) early cardiac and mesoderm genes *T, Eomes, Mesp1, Mixl1, Smarcd3*, (B) structural cardiomyocyte genes *Myl7, Tnnt2, Actc1*, and (C) cardiac transcription factors *Tbx5, Hand1, Nkx2-5, Gata5*.

**Fig. S4.**
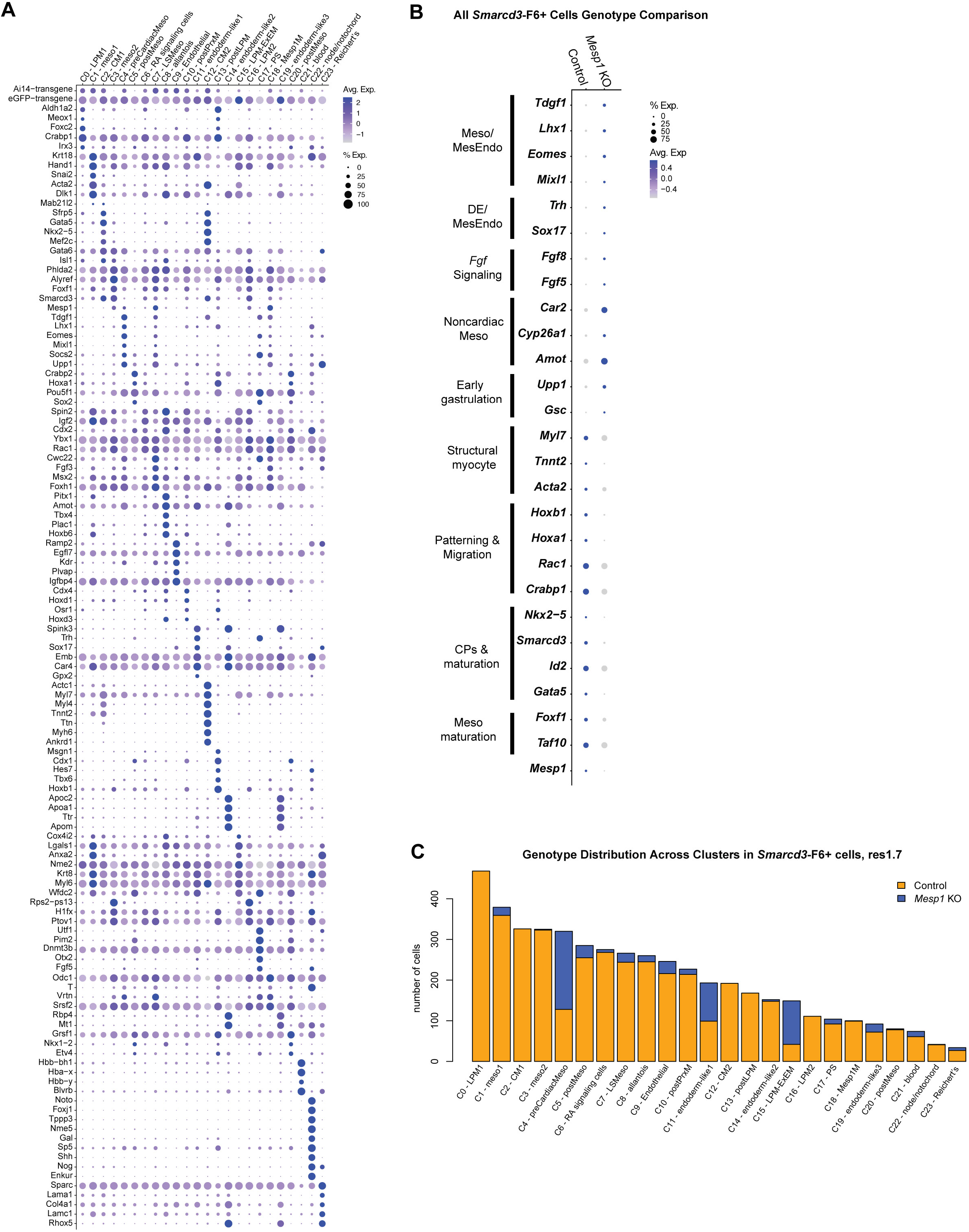
Transcriptional profiles of *Smarcd3*-F6+ cells. (A) Dotplot denoting marker genes and cell type annotations by cluster in *Smarcd3*-F6+ cells atlas. (B) Dotplot representation of differential gene expression between genotypes across all cells. Size of dot denotes percent of cells expressing gene, color of dot represents average gene expression. (C) Barplot denoting distribution of number of cells from genotypes across cluster identities for *Smarcd3*-F6+ atlas.

**Fig. S5.**
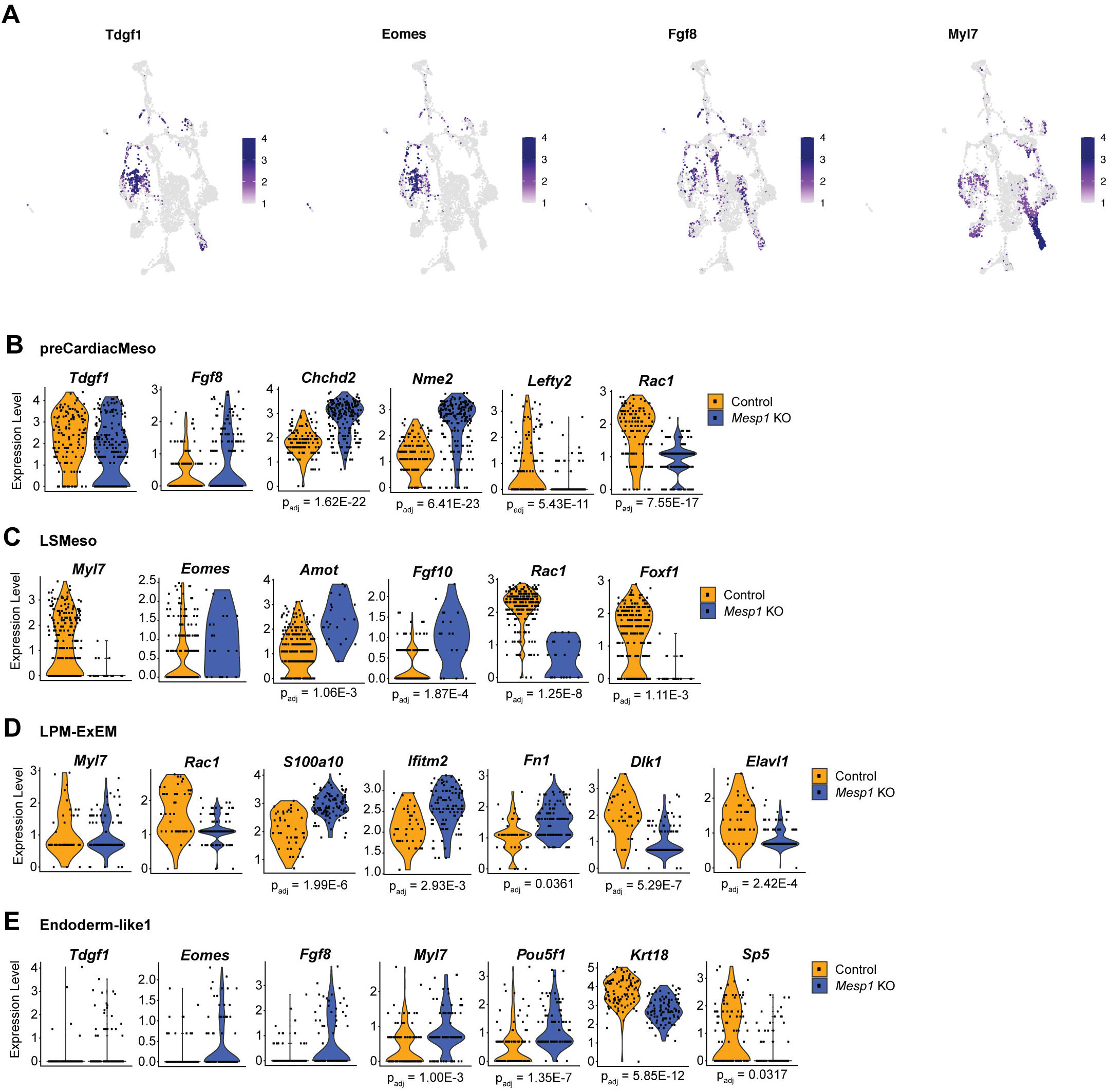
Differentially expressed genes in *Smarcd3*-F6+ cells from *Mesp1* KO embryos. (A) Overlay of gene expression in UMAP space for early cardiac marker genes *Tdgf1*, *Eomes*, *Fgf8*, *Myl7*. (B-E) Differential gene expression profiling highlights similar cardiac marker gene expression between genotypes in (B) preCardiacMeso, (C) LSMeso, (D)LPM-ExEM, and (E) endoderm-like1 cells. (B-E) Differentially expressed genes plotted with adj p values < 0.05.

**Fig. S6.**
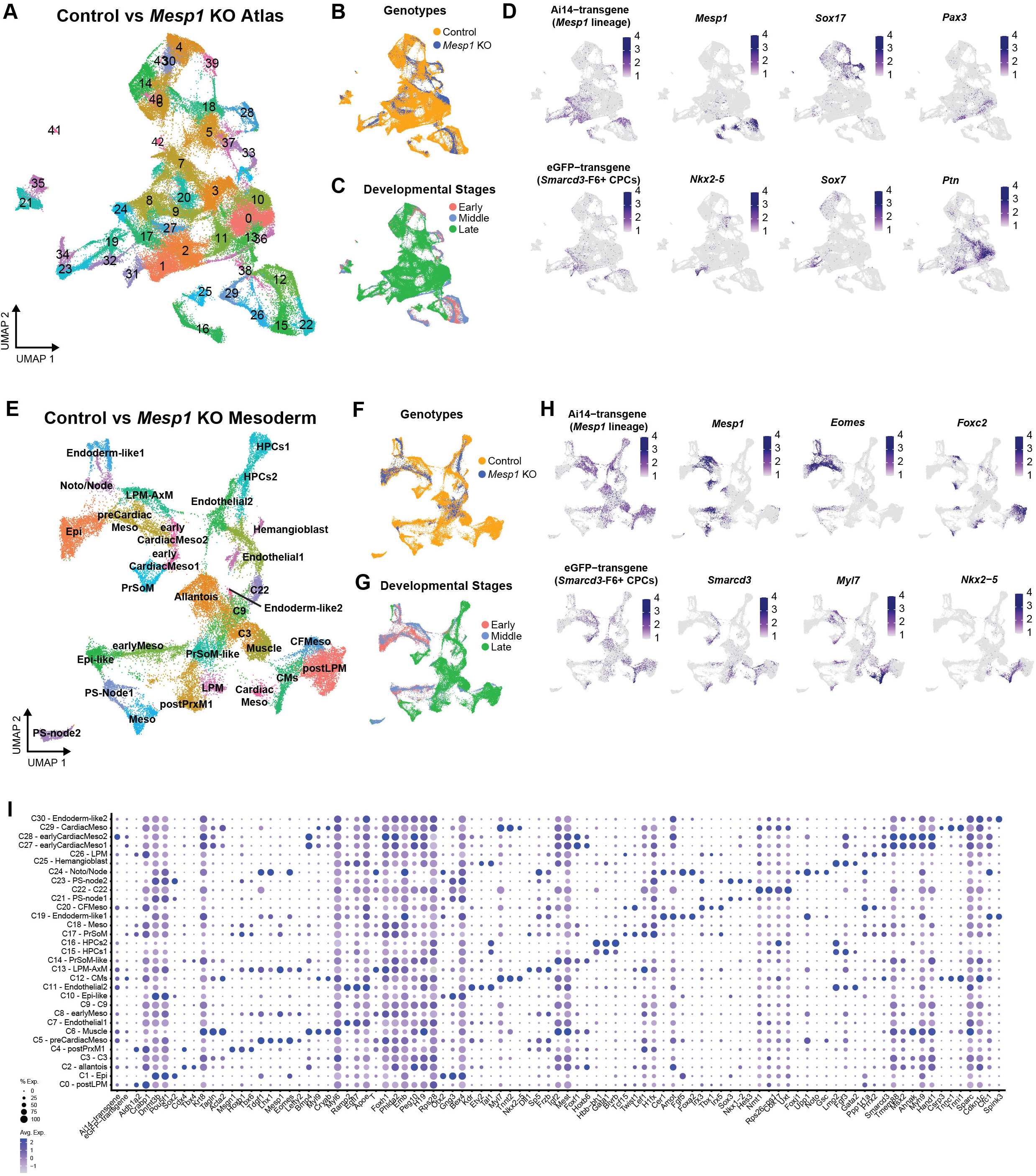
Identification of emerging cardiac mesoderm in control and *Mesp1* KO embryo scRNA-seq data. (A) Atlas UMAP of 96,027 cells representing whole embryos with overlay of (B) genotypes (C) relative developmental stages Early, Middle, and Late. (D) UMAPs showing expression of *Mesp1* lineage transgene Ai14, CPC-specific *Smarcd3*-F6 transgene eGFP, cardiac mesoderm markers *Mesp1*, *Nkx2-5,* endoderm markers *Sox17*, S*ox7*, neural markers *Pax3*, *Ptn*. (E) Atlas UMAP of 35,792 mesoderm cells with overlay of (F) genotypes and (E) relative developmental stages. (H) UMAPs showing gene expression of cardiac and mesoderm genes. (I) Doplot denoting marker genes and cell type annotations by cluster in mesoderm atlas. Size of dot represents percent of cells expressing gene and color represents average expression level. Cluster number used to denote cell types when annotation was not possible.

**Fig. S7.**
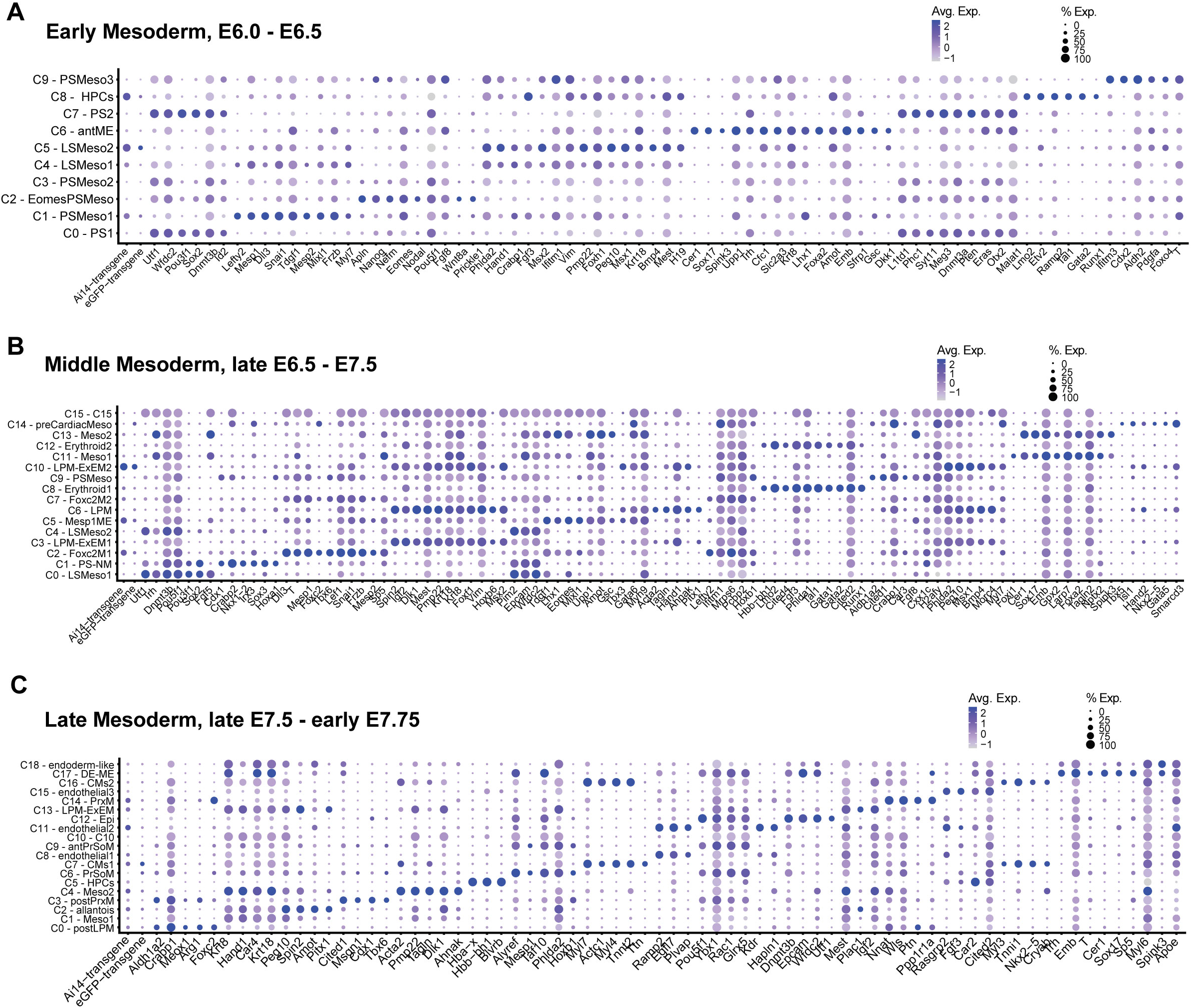
Cell type labels in mesoderm developmental stages atlases. (A) Doplot denoting marker genes and cell type annotations by cluster in Early mesoderm atlas. (B) Doplot denoting marker genes and cell type annotations by cluster in Middle mesoderm atlas. (C) Doplot denoting marker genes and cell type annotations by cluster in Late mesoderm atlas. Size of dot represents percent of cells expressing gene and color represents average expression level.

**Fig. S8.**
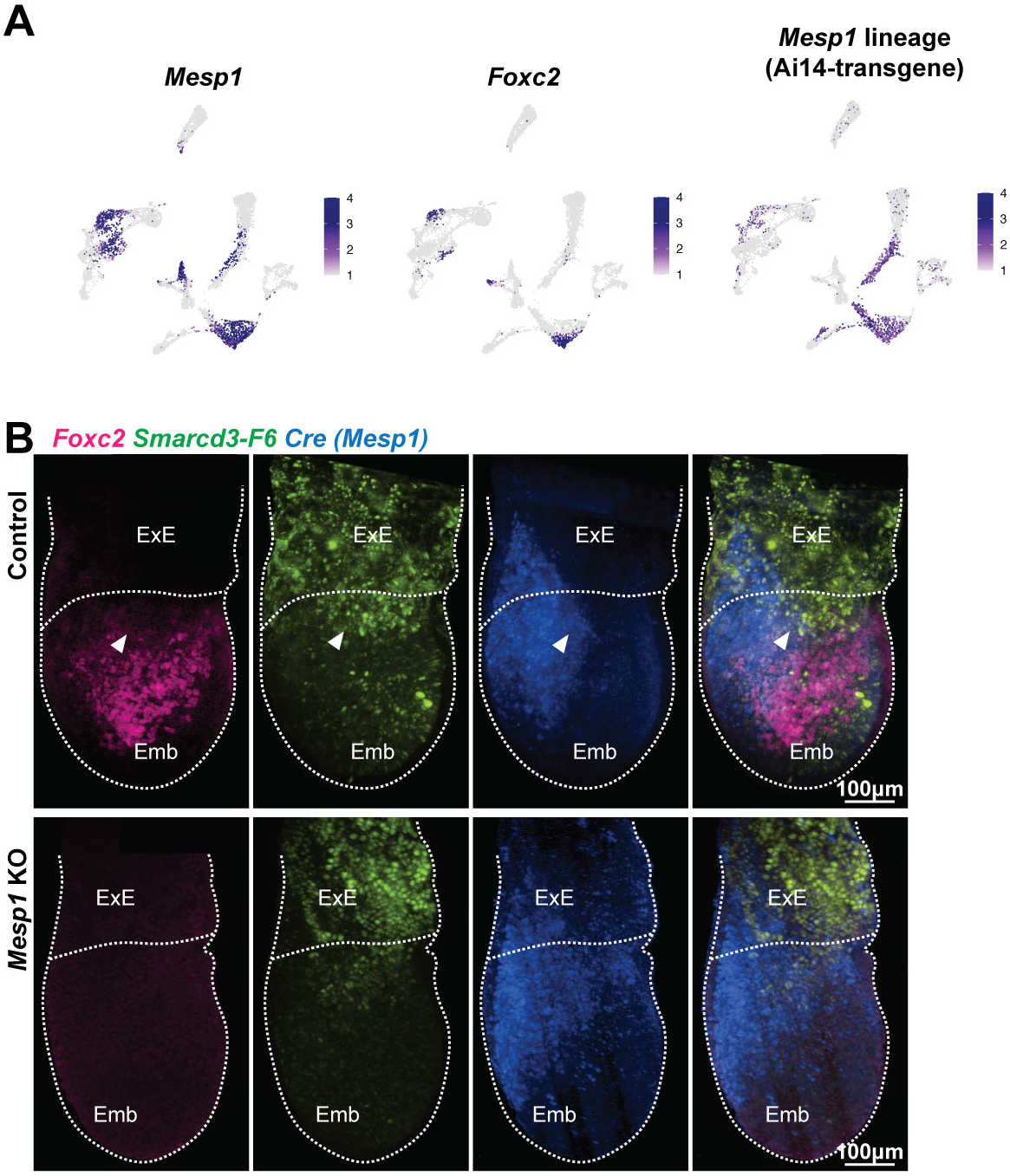
Disrupted organization of mesoderm in Middle stage *Mesp1* KO embryos. (A) Overlay of *Mesp1*, *Foxc2*, and Ai14 *Mesp1*-lineage gene expression in cell types of Middle mesoderm atlas UMAP. (B) Immunostaining and Light Sheet Confocal microscopy for *Foxc2* (magenta), *Smarcd3*-F6 (green) and *Mesp1* via Cre detection (blue) in Middle stage embryos (∼E6.75). Arrowheads denote domain boundaries in control and disruption in *Mesp1* KO embryo. Scale bars are 100 μm.

**Fig. S9.**
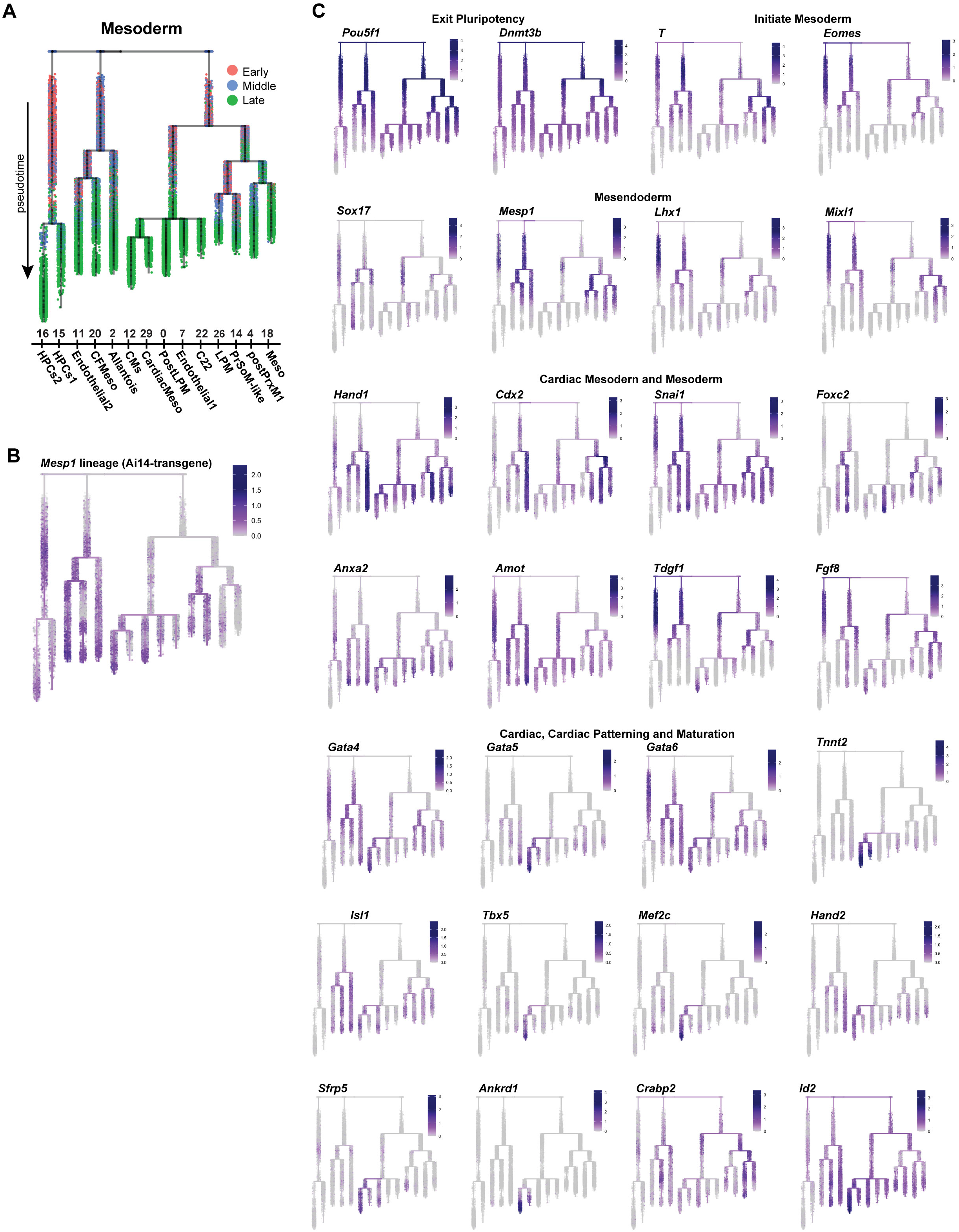
URD trajectory for pseudotime ordering of control and *Mesp1* KO mesoderm. (A) URD tree labeled with relative developmental stages of embryos. (B) URD tree labeled *with Mesp1* lineage transgene reporter Ai14. (C) URD trees labeled with gene expression of various mesodermal genes and TFs involved in regulatory cardiogenesis.

**Fig. S10.**
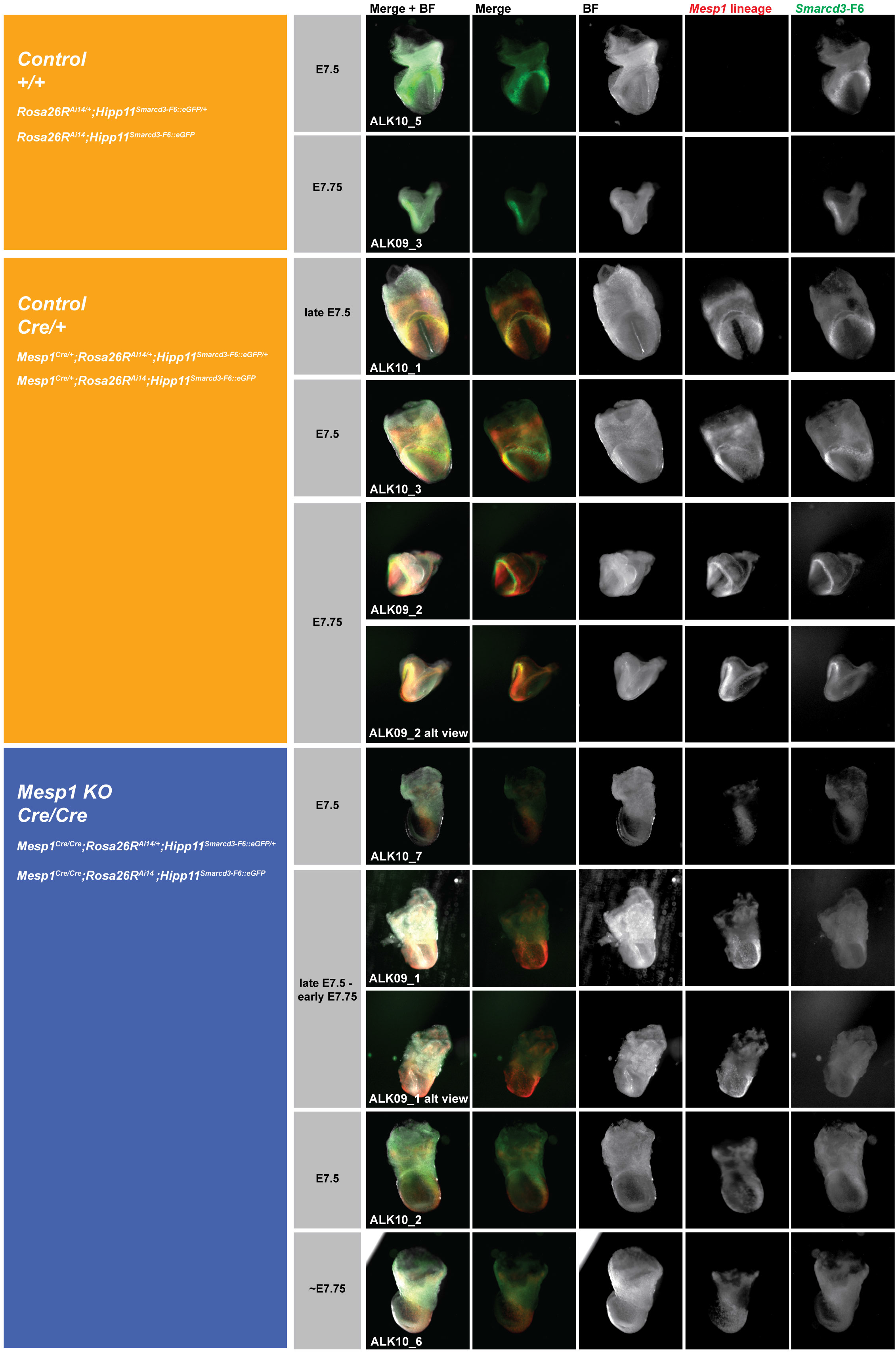
Middle- and Late-stage embryos assayed for scATAC-seq. Images of embryos utilized in generation of control and *Mesp1* KO scATAC-seq dataset. *Mesp1* lineage visualized by endogenous Ai14 fluorescent reporter transgene. *Smarcd3*-F6 visualized by endogenous eGFP fluorescent reporter transgene. Images not acquired and processed identically.

**Fig. S11.**
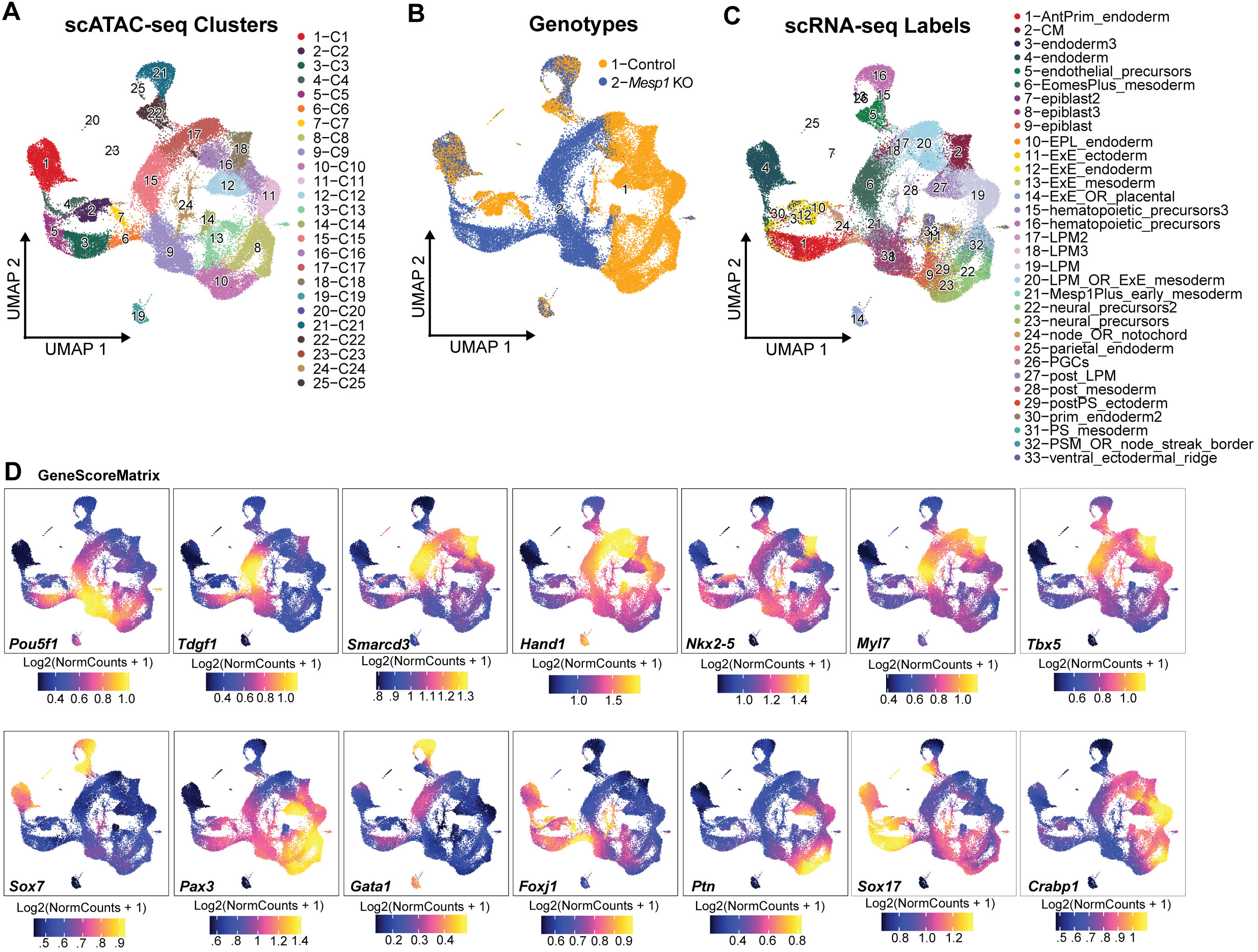
Identification of mesoderm in scATAC-seq data from whole embryos. (A) Whole embryo scATAC-seq atlas with overlays for (B) genotype and (C) relative cell type identities from integration of the complementary scRNA-seq dataset. (D) GeneScoreMatrix plots for chromatin accessibility around gene loci of various mesoderm, cardiac, endoderm, ectoderm, neuronal marker genes.

**Fig. S12.**
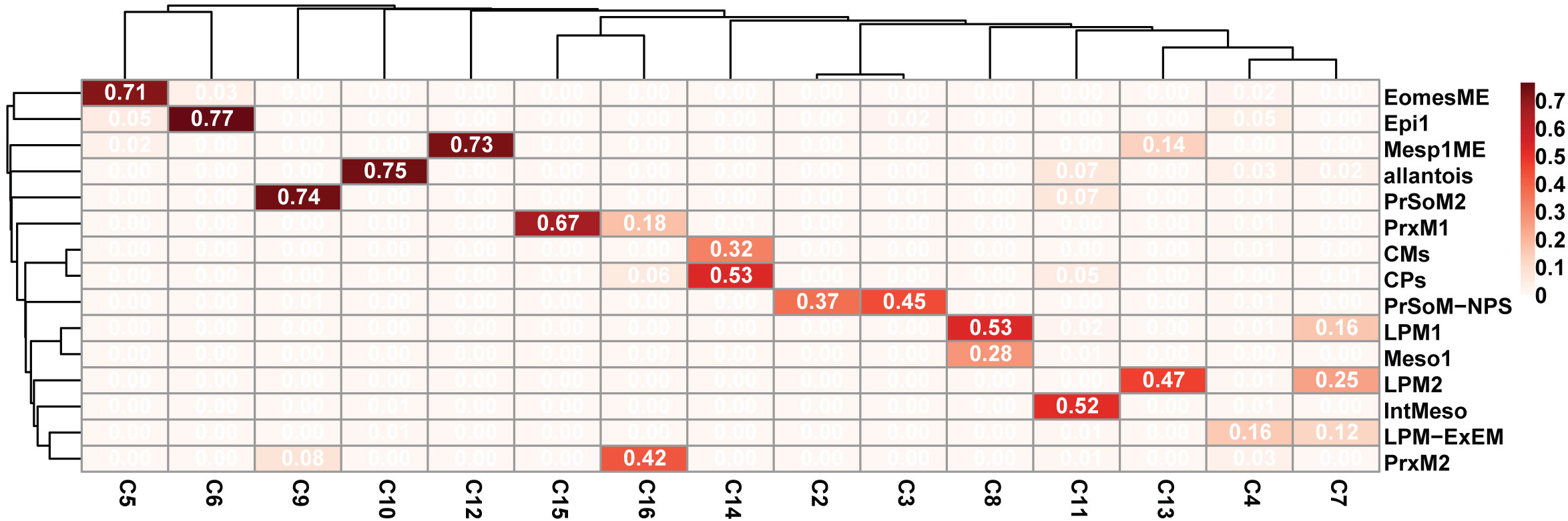
Jaccard Similarity Index for scATAC-seq cluster annotation. Scaled strength of similarity match for scRNA-seq complementary dataset label transfer (rows) onto scATAC-seq clusters (columns). Values 0-1 indicate strength of similarity match for relative cell type annotations. Greater values indicate stronger label matching.

**Fig. S13.**
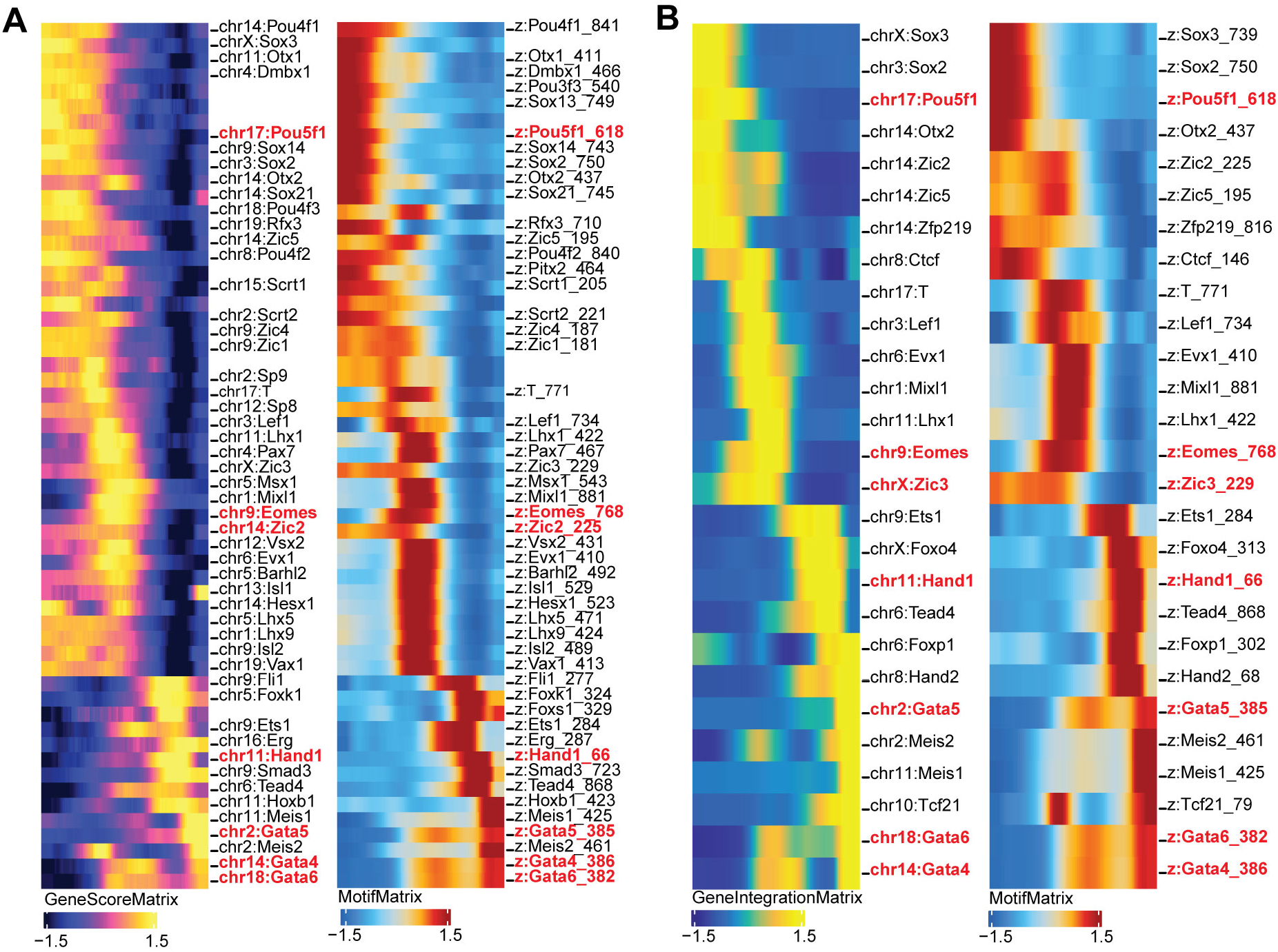
Integrative scATAC-seq trajectory pseudotime correlation analysis. Heatmap visualizations of dynamic shifts along pseudotime progress for correlation matrices (A) between accessibility near TF loci, GeneScoreMatrix, with associated TF motifs, MotifMatrix and (B) between TF gene expression, GeneIntegrationMatrix, with associated TF motifs, MotifMatrix. Motifs in red represent selected putative positive regulators.

**Fig. S14.**
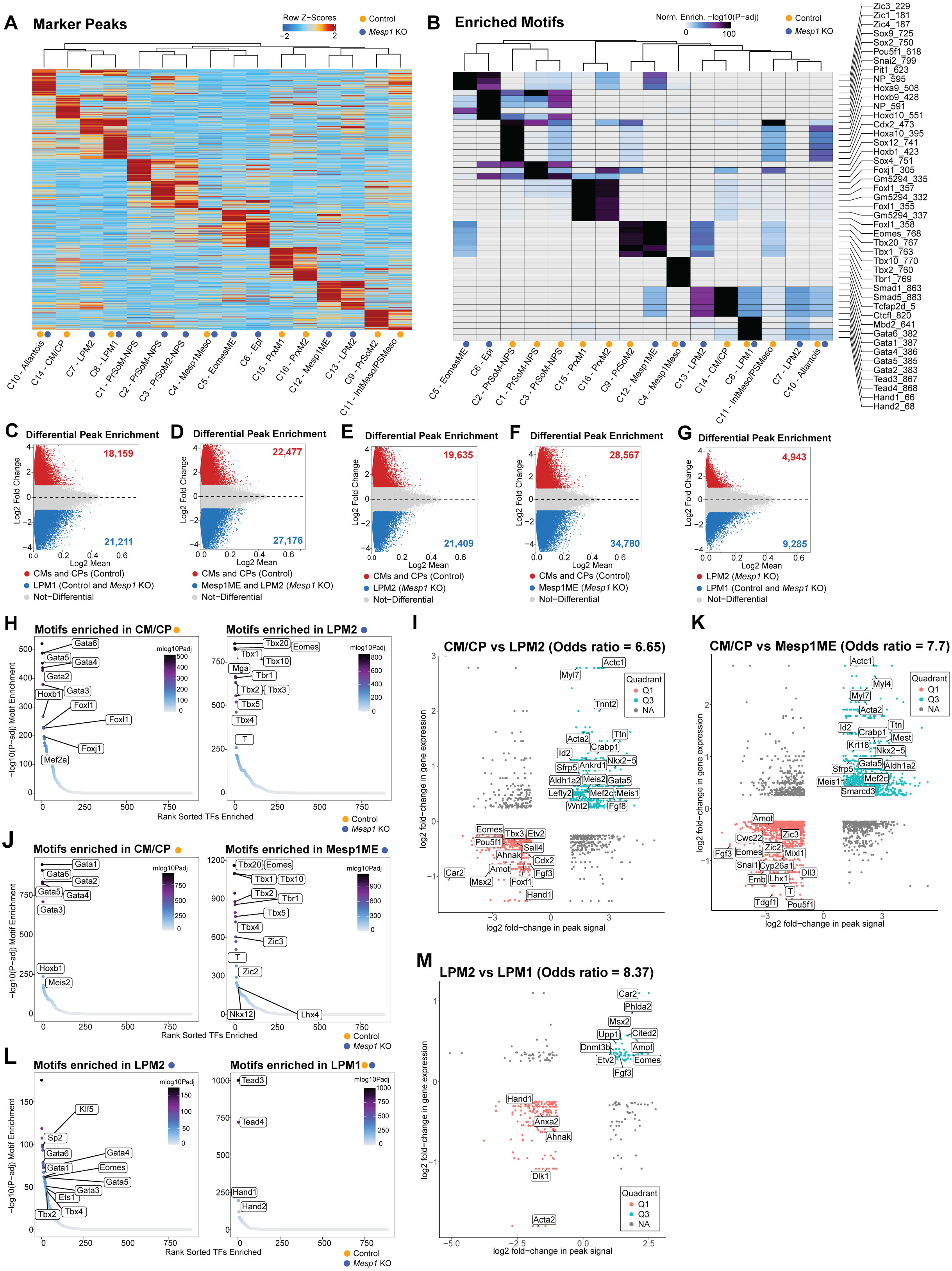
Differential peak and motif enrichment in cardiogenic cell types of control and *Mesp1* KO mesoderm cells. (A) Heatmap for Marker Peak (FDR <= 0.05, Log2FC >=1) accessibility profiles of mesoderm cell types comprised of control, *Mesp1* KO, or both genotypes. (B) Heatmap for enriched motifs (FDR <=0.05, Log2FC >=1) in cluster Marker Peaks. (C-G) MA plots for pairwise comparisons of differential peak enrichment between cardiogenic cell types with genotypes noted. (H, J, L) Motifs enriched in differentially accessible peaks between noted cell types and cluster genotypes. (I, K, M) Plots for peak,gene associations showing correlations between differential peak accessibility and gene expression in comparisons between cells type1 vs type2. Q3 peak,gene pairs represent significantly more accessible peaks paired with upregulated gene expression in type1 cells. Q1 peak,gene pairs represent significantly more accessible peaks paired with upregulated gene expression in type2 cells. Odds ratio denotes probability for observed peak,gene relationships. (I) Peak,gene association plot for cells in CMCP vs LPM2 comparison, (K) CMCP vs Mesp1ME comparison, (M) LPM2 vs LPM1 comparison.

**Fig. S15.**
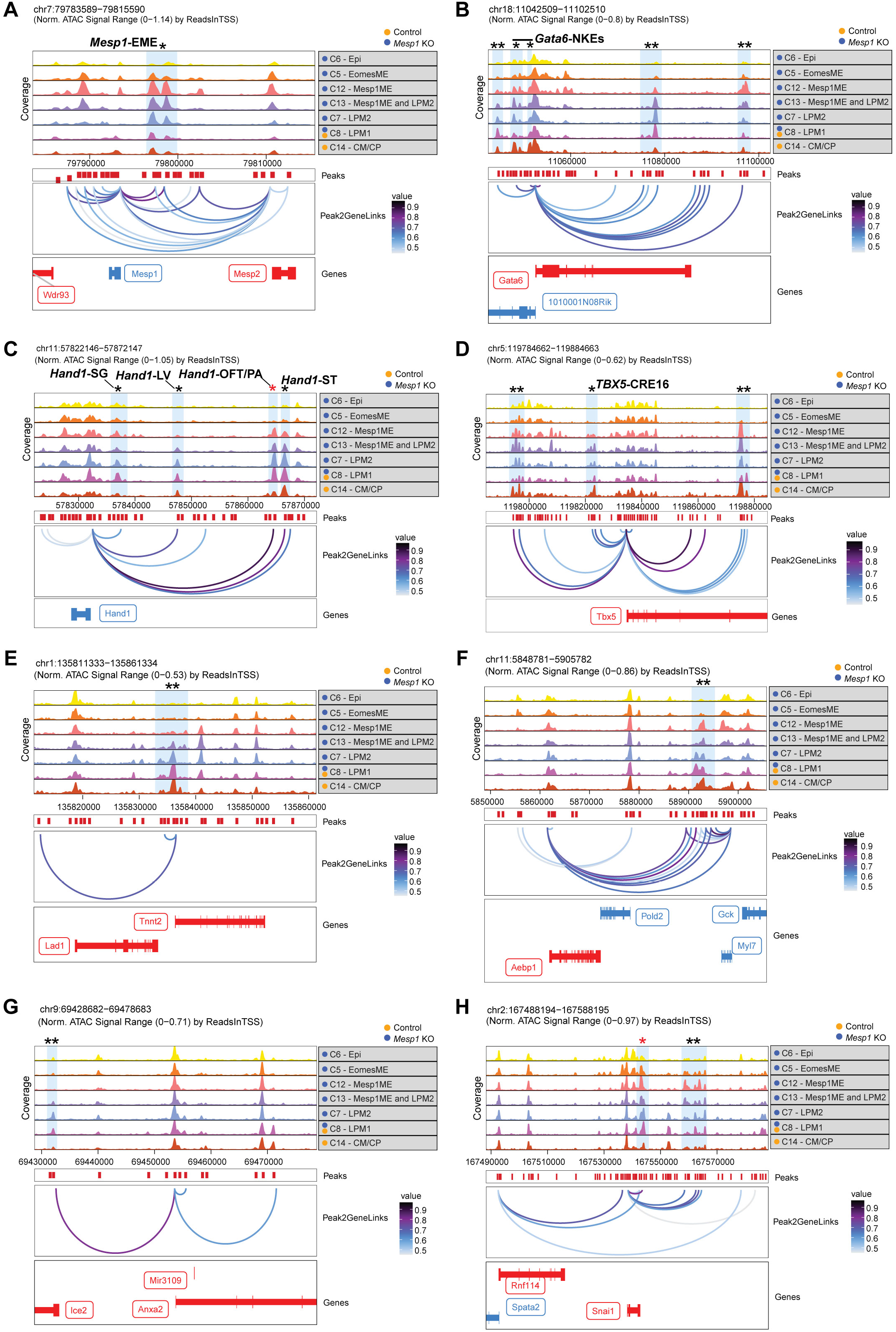
Peak2Gene linkage plots for dysregulated genes in *Mesp1* KO embryos. Peak2Gene linkage browser tracks for cell types showing predicted regulatory connections between distal accessible regions (Peaks) and nearby genes. Shaded bars denote predicted distal regulatory regions; *denotes characterized elements; red* denotes regions with Mesp1-binding; **denotes uncharacterized elements. Characterized elements named when available. Peak linkages to genes (A) *Mesp1* (B) *Gata6,* (C) *Hand1,* (D) *Tbx5,* (E) *Tnnt2*, (F) *Myl7*, (G) *Anxa2*, (H) *Snai1*.

## SUPPLEMENTAL DATA TABLES

**Supplemental Table 1.** Key abbreviations used in this paper.

**Supplemental Table 2.** Corresponds to Fig. 1.

**Supplemental Table 3.** Corresponds to Fig. 2.

**Supplemental Table 4.** Corresponds to Fig. 3.

**Supplemental Table 5.** Corresponds to Fig. 4.

**Supplemental Table 6.** Corresponds to Fig. 5.

**Supplemental Table 7.** Corresponds to Fig. 6.

## MATERIALS AND METHODS

### Mouse models

Animal studies were performed in strict compliance with the UCSF Institutional Animal Care and Use Committee. Mice were housed in a standard 12 hour light/dark animal husbandry barrier facility at the Gladstone Institutes. The *Mesp1^Cre/+^* knock-in mice were obtained from Yumiko Saga (Ajima et al., 2021; Saga et al., 1999). *Rosa26R^Ai14^* mice were from Jackson Laboratory (strain #007914, (Madisen et al., 2010). Tdgf1::*LacZ* mice containing a transgene for *Tdgf1* enhancer with a LacZ reporter were obtained from Brian Black (Barnes et al., 2016).

Control embryos were generated from crosses of *Mesp1^Cre/+^;Rosa26R^Ai14^;Hipp11^Smarcd3-F6::eGFP^* males to C57BL/6J wildtype, *Mesp1^Cre/+^*, or *Mesp1^Cre/+^;Rosa26R^Ai14^;Hipp11^Smarcd3-F6::eGFP^* females. *Mesp1* KO embryos were generated from crosses of *Mesp1^Cre/+^;Rosa26R^Ai14^;Hipp11^Smarcd3-F6::eGFP^* males to *Mesp1^Cre/+^*, or *Mesp1^Cre/+^;Rosa26R^Ai14^;Hipp11 ^Smarcd3-F6::eGFP^* females. Transgenic embryos for single cell transcriptomic and epigenomic sequencing experiments were all on a C57BL/6J background. Transgenic embryos for whole mount *in situ* hybridizations and immunohistochemistry validations were on C57BL/6J backgrounds or a mixed CD1 / C57BL/6J background, in order to facilitate better littermate stage matching via larger litters, with litters born to *Mesp1^Cre/+^* CD1 / C57BL/6J hybrid females mated to *Mesp1^Cre/+^;Rosa26R^Ai14^;Hipp11^Smarcd3-F6::eGFP^* C57BL/6J males. “Control” denotes embryos with at least one wildtype allele in the *Mesp1* locus and includes genotypes *Mesp1^Cre/+^;Rosa26R^Ai14^;Hipp11^Smarcd3-F6::eGFP^*, *Mesp1^Cre/+^;Rosa26R^Ai14/+^;Hipp11^Smarcd3-F6::eGFP/+^*, *Mesp1^+/+^;Rosa26R^Ai14^;Hipp11^Smarcd3-F6::eGFP^,* or *Mesp1^+/+^;Rosa26R^Ai14/+^ Hipp11^Smarcd3-F6::eGFP/+^.* Heterozygosity of *Mesp1^Cre/+^* or *Mesp1^+/+^* is noted when control embryos were utilized in scRNA-seq (Fig. S1) or scATAC-seq (Fig. S10) library generation. “*Mesp1* KO” denotes embryos with homozygosity of the Cre insertion disrupting the *Mesp1* locus and includes genotypes *Mesp1^Cre/Cre^;Rosa26R^Ai14^;Hipp11^Smarcd3-F6::eGFP^* or *Mesp1^Cre/Cre^;Rosa26R^Ai14/+^;Hipp11^Smarcd3-F6::eGFP/+^*.

Control embryos for activity assessment of the *Tdgf1* enhancer had genotypes *Mesp1^Cre/+^;Tdgf1*::LacZ or *Tdgf1*::LacZ, and *Mesp1* KO embryos had genotypes *Mesp1^Cre/Cre^:Tdgf1*::LacZ.

### Cloning and generation of TARGATT transgenic knock-in mice

The *Smarcd3*-F6 fragment was isolated and cloned with inclusion of an *nlsEGFP* under control of an *Hsp68* minimal promoter for TARGATT (Applied Stem Cells) insertion to the *Hipp11* locus as previously described (Devine et al., 2014) to create the *Hipp11^Smarcd3-F6::eGFP^* mouse. Purified construct DNA was injected into embryo pronuclei along with mRNA for the *Phi31o* transposase according to manufacturer’s protocols.

### Timed matings and whole embryo dissections

To achieve timed matings, male and female mice were housed together in the evening and pregnancy was assessed by vaginal plug the following morning. Gestational stage was determined starting as day E0.5 at noon of plug detection. Females were confirmed pregnant by abdominal ultrasound (Vevo 3100, Visual Sonics) the afternoon of day 6 or morning of day 7 and sacrificed according to IACUC standard procedure at noon on day 7, or the early morning of day 8. The embryonic ages captured in individual litters ranged from E6.0 to E7.5 on day 7, and E7.5 to E7.75 on day 8. The diversity of ages in litters aided in the construction of a fine timecourse for both mutant and control timelines.

Embryos were dissected and in later stages when yolk is present, also de-yolked, in ice-cold PBS (Life Technologies, 14190250) with 1% FBS (Thermo Fisher Scientific, 10439016) on ice. Embryos were screened using an upright epifluorescent dissecting microscope (Leica MZFLIII microscope, Lumen Dynamics XCite 120LED light source, Leica DFC 3000G camera) for presence of both red and green fluorescent reporters, indicative of *Mesp1* lineage tracing from *Mesp1^Cre^*;*Rosa26R^Ai14^* alleles and expression of the *Smarcd3*-F6::eGFP transgene reporter from the *Hipp11^Smarcd3-F6::eGFP^* allele, respectively. Embryos were staged according to (Downs & Davies, 1993). For difficult-to-capture control stages used in construction of the wildtype scRNA-seq timeline, absence of *Mesp1* lineage (Ai14) reporter was permitted and noted for those embryos (Fig. S1). Additionally, *Mesp1^+/+^* alleles were specifically included in addition to *Mesp1^Cre/+^ as* controls for scATAC-seq library generation in the event locus-specific effects of Cre insertion required additional consideration, which we didn’t find to be the case as both control genotypes appeared identically in the dataset (Fig. S10). DNA for genotyping was extracted using QuickExtract DNA Extraction Solution (Lucigen, QE09050) from harvested yolk sac tissue if available or else from a micro-dissected nick of the extraembryonic anterior proximal region. Genotyping was performed to distinguish *Mesp1* KO embryos from control embryos using Phire Green Hot Start II DNA Polymerase (Thermo Fisher Scientific, F124L) according to manufacturer’s protocols using primers to detect wildtype bands (control, P1+P3) and Cre alleles (*Mesp1* KO, P1+P2):

*Mesp1* FWD, P1: GGC CAT AGG TGC CTG ACT TA
Cre2 REV, P2: CCT GTT TTG CAC GTT CAC GG
*Mesp1* REV, P3: ACC AGC GGG ACT CAG GAT

### Embryo preparation for single-cell library generation

Due to the small size and lack of morphological distinction between tissue types of embryos at these early stages, whole embryos were dissected and harvested for single cell library generation.

Whole embryos were incubated in 200 μL 0.25% TrypLE (ThermoFisher Scientific, 12563029) solution for 5 min at 37°C and triturated gently. Dissociated cell suspension was quenched with 600 μL of PBS with 1% FBS, singularized via passage through a 70 μm cell strainer (BD Falcon, 352235), pelleted by centrifugation at 150xg for 3 min, and resuspended in 34 μL of PBS with 1% FBS. At least 2 embryos were collected per genotype per embryonic stage in all datasets except for the *Mesp1* KO embryos in the scRNA-seq dataset where this was not possible, and the use of relative developmental stages was employed in analysis along with replicate validations via *in-situ* hybridization for differentially expressed genes.

### Single-cell transcriptome library preparation and sequencing

Libraries for scRNA-seq were prepared according to manufacturer’s instructions using the 10X Genomics Chromium controller, Chromium Single Cell 5’ Library and Gel Bead Kit v1 (10X Genomics, 1000006) and Chromium Single Cell A Chip Kit (10X Genomics, 1000151). A maximum of 10,000 cells per sample were loaded onto the 10X Genomics Chromium instrument, and each sample was indexed with a unique sample identifier (10X Genomics Chromium i7 Multiplex Kit, 120262). Final libraries were pooled and sequenced shallowly according to 10X protocol parameters on a NextSeq500 (Illumina), and then re-pooled for deeper sequencing on HighSeq4000 (Illumina) and/or NovaSeq using an S4 lane (Illumina). Littermate, stage-matched comparisons of control and *Mesp1* KO libraries were always sequenced together in the same library pool. All scRNA-seq libraries were sequenced to a mean read depth of at least 50,000 total aligned reads per cell.

### Processing raw scRNA-seq

Raw sequencing reads were processed using the 10X Genomics Cellranger v3.0.2 pipeline. Reads were demultiplexed using cellranger mkfastq and aligned with cellranger count to the Mm10 reference genome containing additional sequences for the Ai14 and eGFP. Cellranger “aggr” was used to aggregate and read depth normalize multiple GEM libraries for either the wildtype atlas dataset or the atlas dataset containing control and *Mesp1* KO embryo libraries.

### Seurat analysis of scRNA-seq data

Outputs from the Cellranger pipeline were analyzed using the Seurat Package v3.0.2 in R (Butler et al., 2018; Stuart et al., 2019; Satija et al., 2015). The dataset containing all wildtype embryos and the “WTvsMut” dataset containing control (wildtype) and *Mesp1* KO embryos were analyzed as separate Seurat objects. A single aggregated counts matrix for each separate dataset were used as inputs for Read10X and CreateSeuratObject functions. Quality control steps were performed to remove dead cells or doublets.

#### Wildtype Atlas

For the wildtype atlas, cells with <10% mitochondrial reads, UMI counts less than 50,000, and detected genes between 200 and 6,300 were retained. SCTransform (Hafemeister & Satija, 2019) was used to normalize and scale data with regressions performed with respect to mitochondrial percent, number of genes, and number of UMI counts detected. PCA analysis and batch correction were performed using FastMNN (Haghverdi et al., 2018) split by experimental group (experiment number denoted with library prefixes ALK06, ALK08, ALK07, ALK05, ALK04). 94,824 cells were clustered based on the top 50 principal components and visualized using RunUMAP, FindNeighbors, and FindClusters and outputs were visualized as Uniform Manifold Approximation and Projection (UMAP) embeddings generated with DimPlot. Cell types were annotated at clustering resolution 0.4 using the FindAllMarkers function with Wilcoxon rank-sum test (min.pct = 0.1, logfc threshold = 0.25) to identify cluster specific marker genes. Relevant mesoderm cell types were subsetted based on cluster-wise detection of *Smarcd3*-F6::eGFP and Ai14 transgenes for CPCs and the *Mesp1* lineage, respectively. The resulting 34,724 were re-clustered and re-annotated at resolution 1.2 to create the cardiac mesoderm wildtype atlas.

#### Whole Embryo Control vs. Mesp1 KO Atlas

For the WTvsMut atlas, cells with <10% mitochondrial reads, UMI counts less than 50,000, and detected genes between 200 and 7,000 were retained. SCTransform was used to normalize and scale data with regressions performed with respect to mitochondrial percent, number of genes, and number of UMI counts detected. PCA analysis and batch correction were performed using FastMNN split by experimental group as in wildtype dataset. Cells were clustered as described for wildtype atlas above, with iterative clustering performed following removal of low quality clusters. This WTvsMut dataset represents 96,027 cells containing 79,725 control and 16,302 *Mesp1* KO cells. Cluster cell types were annotated at resolution 1.0 using FindAllMarkers as described above.

The relevant developmental stages were annotated within Seurat meta data. Cells from 6 embryos staged E6.0 - E6.5 (ALK06_2_E60_con_rep1, ALK06_4_E60_con_rep2, ALK08_20_E60_con_rep3, ALK08_14_lateE60_con_rep1, ALK07_15_E65_con_rep1, ALK08_6_E65_Mesp1KO_rep1) were denoted as “Early” stages. Cells from 4 embryos staged late E6.5 – early E7.5 (ALK07_3_lateE65_con_rep1, ALK07_14_E70_con_rep1, ALK08_11_E70_Mesp1KO_rep1, ALK07_7_earlyE75_con_rep1) were denoted as “Middle” stages. Cells from 5 embryos staged late E7.5 to early E7.75 when cardiac crescent is formed (ALK07_6_lateE75_con_rep1, ALK04_3_lateE75_con_rep2, ALK05_7_E775_con_rep1, ALK05_2_lateE75_Mesp1KO_rep1, ALK07_8_E775_con_rep2) were denoted as “Late” stages. While we set out to acquire replicates of both genotypes per each stage as the most optimal statistical scenario, the 25% yield of *Mesp1* KO embryos within C57BL/6J litter sizes at these early gastrulation stages proved prohibitive. Thus we relied on validations of key scRNA-seq findings via the orthogonal approach of multiplexed whole mount *in situ* hybridizations.

#### Smarcd3-F6+ Control vs Mesp1 KO Atlas

To analyze putative CPCs, all cells expressing the *Smarcd3-*F6-eGFP transgene were subsetted from the full WTvsMut atlas and re-clustered into their own Seurat object containing 4,868 cells (4,276 control and 592 *Mesp1* KO cells). FindAllMarkers function was used to identify cluster marker genes of represented cell types at resolution 1.7. The analysis between control and *Mesp1* KO genotypes irrespective of cell type was performed using FindMarkers function between genotypes with Wilcoxon rank-sum test (min.pct = 0.1, logfc threshold = 0.25). Cluster-wise differential gene expression testing was performed using FindMarkers function and Wilcoxon rank-sum test (min.pct = 0.1, logfc threshold = 0.25) between genotypes within specific cell type clusters, and visualized with the VlnPlot function. Differential gene expression results irrespective of cell type were visualized by DotPlot function separated by genotypes and also genotypes separated by developmental stages.

#### Mesoderm Control vs Mesp1 KO Atlas

Relevant mesoderm cells were subsetted from the full WTvsMut atlas based on cluster-wise detection via FeaturePlot and VlnPlot at cluster resolution 1.0 of *Smarcd3*-F6::eGFP and Ai14 transgenes for CPCs and the *Mesp1* lineage, respectively. The resulting 35,792 cells (29,924 control and 5,868 *Mesp1* KO cells) of the WTvsMut mesoderm dataset was re-clustered and annotated at resolution 1.5 using FindAllMarkers function as above to identify cell type marker genes as described above.

Embryos representing the relative developmental stages of “Early” (5,504 cells; 4,472 control and 1,032 *Mesp1* KO), “Middle” (7,666 cells; 6,734 control and 932 *Mesp1* KO), and “Late” (22,622 cells; 18,718 control and 3,904 *Mesp1* KO) as described above were subsetted into respective individual Seurat objects, re-clustered as described, and cell type clusters were further re-annotated (at resolutions 0.7, 0.7, 0.7 for Early, Middle, and Late objects, respectively). Clusters representing cell types relevant for cardiac development were identified through cluster-wise enrichment of *Smarcd3*-F6::eGFP transgene expression overlayed in UMAP space via FeaturePlot. Differential gene expression testing between genotypes within cardiogenic cell type clusters was performed using FindMarkers function with Wilcoxon rank-sum test (min.pct = 0.1, logfc threshold = 0.25). Differentially expressed genes with adjusted p-values < 0.05 were plotted as violin plots in Seurat except in cases to highlight total absence of transcript in one genotype condition.

#### Whole Embryo Control vs Mesp1 KO Atlas for scATAC-seq integration

For the scRNA-seq WTvsMut atlas for integration with scATAC-seq data, libraries from Middle stage embryos (ALK07_3_lateE65_con_rep1, ALK07_14_E70_con_rep1, ALK08_11_E70_Mesp1KO_rep1, ALK07_7_earlyE75_con_rep1) and Late stage embryos (ALK07_6_lateE75_con_rep1, ALK04_3_lateE75_con_rep2, ALK05_7_E775_con_rep1, ALK05_2_lateE75_Mesp1KO_rep1, ALK07_8_E775_con_rep2) were subsetted from the aggregated WTvsMut counts matrix. Cells with <7.5% mitochondrial reads, UMI counts less than 50,000, and detected genes between 200 and 7,000 were retained. SCTransform was used to normalize and scale data with regressions performed with respect to mitochondrial percent, number of genes, and number of UMI counts detected. PCA analysis and batch correction were performed using FastMNN split by experimental group. After initial clustering as previously described, cell clusters representing low quality cells were removed and clustering was iterated again. The resulting dataset represents 82,536 cells containing 68,717 control and 13,819 *Mesp1* KO cells. Cluster cell types were annotated at resolution 1.2 using FindAllMarkers as described above.

#### Mesoderm Control vs Mesp1 KO Atlas for scATAC-seq integration

Relevant mesoderm cells were subsetted from the whole embryo matched scATAC-seq WTvsMut atlas based on cluster-wise detection via FeaturePlot and VlnPlot of *Smarcd3*-F6::eGFP and Ai14 transgenes for CPCs and the *Mesp1* lineage, respectively. The resulting 30,427 cells (26,054 control and 4,373 *Mesp1* KO cells) of the scATAC-seq matched mesoderm WTvsMut dataset were re-processed from RNA assay slot with the standard Seurat workflow NormalizeData, FindVariableFeatures and ScaleData. SCTransform was not used in this mesoderm scRNA-seq dataset because we found that while cell type label-transfer with scATACseq was successful as previously described for the whole embryo integration, downstream scATAC-seq analyses leveraging the scRNA-seq gene integration matrix performed in the mesoderm scATAC-seq dataset were incompatible with SCT-normalized values. PCA analysis and batch correction were performed using FastMNN split by experimental group. From here clustering was performed as previously described and cell types were annotated at resolution 1.2 using FindAllMarkers function as above to identify cell type marker genes as described above.

Differential gene expression testing between genotypes within cell type clusters and between cell type clusters was performed using FindMarkers function with Wilcoxon rank-sum test (min.pct = 0.1, logfc threshold = 0.25). These lists of differentially expressed genes served as inputs to the (peak, gene) association analyses with scATAC-seq differential peaks using rGreat (below in methods).

### Single cell transcriptomic cell trajectories and pseudotime analysis

Pseudotime analysis was performed using the URD package (version 1.0.2 and 1.1.1) (Farrell et al., 2018). The WTvsMut mesoderm Seurat object containing all three relative developmental stages, processed as previously described, was converted to an URD object using the seuratToURD function. Cell-to-cell transition probabilities were constructed by setting the number of near neighbors (knn) to 189 and sigma to 10. Pseudotime was then calculated by running 80 flood simulations with *Pou5f1*+ epiblast cells containing “Early” staged embryos (cluster 1 of WTvsMut mesoderm Seurat object at resolution 1.5) as the “root” cells. Clusters containing the most defined mesodermal derivative cell types and containing the “Late” staged embryos were set as the “tip” cells (C15-,C16-HPCs, C11-, C7-Endothelial, C20-CFMeso, C2-Allantois, C12-CMs, C29-CardiacMeso, C0-postLPM, C22, C26-LPM, C14-PrSoM-like, C4-postPrxM1,C18-Meso). The resulting URD tree was subsequently built by simulated random walks from each tip. Overlay of relative developmental stages from embryo data was used to show consensus in pseudotime estimations of cell trajectories. Overlay of *Smarcd3*-F6::eGFP and various cardiac marker genes such as *Nkx2-5*, *Myl7*, *Smarcd3*, *Tnnt2*, and various *Gata* transcription factors were used to identify the relevant cardiac-fated branching segments of the URD tree.

To identify differentially expressed genes in fate-related cells of the cardiac branches, cell barcodes from relevant branch segments were extracted from the URD object and assigned their relevant segment branch identities in the corresponding Seurat object. Differential gene testing using the Wilcoxon rank sum test (min.pct = 0.1, logfc threshold = 0.25) was then performed between genotypes within a segment or between noted segments related in their pseudotemporal progression. Differentially expressed genes with adjusted p-values less than 0.05 were plotted as violin plots in Seurat and representative genes were overlayed on the URD tree to visualize expression patterns in pseudotime space.

### Single cell Assay for Transposase Accessible Chromatin (scATAC-seq) library generation

For scATACseq library generation we used the 10X Genomics Chromium, scATACseq library kit v1 (10X Genomics, 1000110) and Chromium Chip E (10X Genomics, 1000156) according to manufacturer’s protocols. Embryos were dissected and dissociated into single cells as described above and cells were resuspended in pre-chilled Lysis buffer for isolation of single nuclei. A maximum of 10,000 nuclei per sample were subjected to transposition and loaded into the 10X Genomics Chromium instrument. Final libraries were pooled and sequenced shallowly according to 10X protocol parameters on a NextSeq500 (Illumina). Littermate, stage-matched comparisons comprising a total of 5 control and 4 *Mesp1* KO embryos were ultimately re-pooled and sequenced together for deep sequencing on a NovaSeq6000 S4 lane (Illumina). All libraries were sequenced to depths of at least 24,000 median fragments per cell, and at most 35,000 median fragments per cell.

### Processing raw scATAC-seq

Raw sequencing reads were processed using the 10X Cellranger ATAC v1.2.0 software pipeline. Reads were demultiplexed using cellranger-atac mkfastq. Cell barcodes were filtered and aligned to the Mm10 reference genome using cellranger-atac count. The resulting output indexed fragment files from each library were not aggregated and served as the inputs for downstream computational analysis in ArchR (Granja et al., 2021).

### ArchR analysis of scATAC-seq

Downstream computational analysis of scATAC-seq data was done with the ArchR software package v1.0.1 in R (Granja et al., 2021). Initial Arrow files were generated for all samples from inputs of respective indexed fragment files and sample meta-data. Samples from embryos aged E7.5 were called “Middle” stage (libraries ALK10_5_E75_con_rep1, ALK10_3_E75_con_rep2, ALK10_1_lateE75_con_rep1, ALK10_7_E75_Mesp1KO_rep1, ALK10_2_E75_Mesp1KO_rep2). Samples from embryos aged E7.75 were called “Late” stage (libraries ALK09_3_E775_con_rep1, ALK09_2_E775_con_rep2, ALK09_1_E775_Mesp1KO_rep1, ALK10_6_E775_Mesp1KO_rep2). The function createArrowFiles was run on each sample, removing cells with a transcription start site (TSS) enrichment score less than 4, and fragments less than 5000. This initialization also creates a genome-wide TileMatrix of 500 base pair bins and a weighted calculation of accessibility within and surrounding gene loci annotated from the Mm10 genome, called a GeneScoreMatrix. While CellRanger v1.2.0 implements removal of multi-cell capture, ArchR recommends an additional round of cell doublet removal using functions addDoubletScores and filterDoublets. Individual ArrowFiles for each sample were aggregated into a single WTvsMut whole embryo ArchRProject containing 46,819 cells (26,295 control, 20,524 *Mesp1* KO) with a median TSS enrichment score of 10.675 and median of 30,703 fragments per cell. Dimensionality reduction was performed with addIterativeLSI (2 iterations, resolution 0.2, 30 dimensions). Clustering was performed using addClusters with “Seurat” method (resolution 0.8) and addUMAP was used to embed values for dimensionality reduced visualizations with the function plotEmbedding. Relative cell-type annotation of clusters was performed with consideration of combined information from GeneScore plots and label transfer from the complementary annotated whole embryo WTvsMut scRNA-seq Seurat analysis object of stage-matched control and *Mesp1* KO embryos for the relative Middle (embryos ALK07_3_lateE65_con_rep1, ALK07_14_E70_con_rep1, ALK08_11_E70_Mesp1KO_rep1, ALK07_7_earlyE75_con_rep1) and Late (embryos ALK07_6_lateE75_con_rep1, ALK04_3_lateE75_con_rep2, ALK05_7_E775_con_rep1, ALK05_2_lateE75_Mesp1KO_rep1, ALK07_8_E775_con_rep2) stages. For scRNA-seq integration, the addGeneIntegrationMatrix function utilizes Seurat’s FindTransferAnchors to perform Canonical Correlation Analysis. Relevant mesoderm clusters (“C15”, “C9”, “C24”, “C17”, “C16”, “C18”, “C12”, “C11”, “C8”) were identified based on relative overlay of scRNA-seq cell type labels onto scATAC-seq clusters and GeneScoreMatrix for key marker genes, and subsetted into a WTvsMut mesoderm ArchRProject containing 25,848 cells (14,212 control and 11,636 *Mesp1* KO).

Dimensionality reduction was performed on the subsetted WTvsMut mesoderm ArchRProject with addIterativeLSI (4 iterations, resolution 0.2, 30 dimensions), which was then batch corrected using addHarmony. Harmonized clustering was then performed using addClusters with “Seurat” method (resolution 0.8) and addUMAP was performed. Clusters were visualized using plotEmbedding. Relative cell-type annotation of clusters was again performed following integration with the mesoderm WTvsMut complementary, annotated, Seurat analysis scRNA-seq object from stage-matched control and *Mesp1* KO embryos for the relative Middle and Late stages. The addGeneIntegrationMatrix function was used to generate GeneIntegration plots, which were compared to GeneScore plots for understanding of cluster markers. A Jaccard Similarity Analysis from the predicted scRNA-seq integration for scATAC-seq clusters annotation was performed similarly to as described (Sarropoulos et al., 2021) to assess the strength of predictive labels, and the resulting proportions were visualized with the pheatmap function from the ComplexHeatmap R package (Gu, Eils & Schlesner, 2016). Cluster identities from the mesoderm subset scATAC-seq dimensionality reduction were utilized for downstream cluster-wise analyses.

#### Peak calling and motif enrichment

Peaks were called using pseudo-bulkification and MACS2. Cell replicates for pseudobulks were created using addGroupCoverages on scATAC-seq clusters (40 minimum and 500 maximum cells in a replicate, minimum 2 replicates per cluster, 0.8 sampling ratio, kmerlength for Tn5 bias correction of 6). Peaks were called using addReproduciblePeakSet (500 peaks per cell, 1.5E5 maximum peaks per cluster) with MACS2 (-75 base pair shift per Tn5 insertion, 150 basepair extension after shift, excluding mitochondrial chromosome genes and chromosome Y genes, with a q-value significance cutoff 0.1). Peaks were then merged using ArchR’s iterative overlap method. Cluster enriched marker peaks were identified with getMarkerFeatures (FDR <= 0.05, Log2FC >=1) and visualized with plotMarkerHeatmap. Cluster motif enrichment was ascertained with addMotifAnnotations using the CIS-BP database motif set. Cluster enriched motifs were visualized with peakAnnoEnrichment (FDR <=0.05, Log2FC >=1) and then the top 7 motifs per cluster were plotted with plotEnrichHeatmap and ComplexHeatmap. Single cell resolution motif enrichment was computed using the chromVAR package (Schep et al., 2017) by adding background peaks (addBgdPeaks) and then motif z-score deviations were computed per cell with addDeviationsMatrix. Motif enrichments were visualized in UMAP embeddings with plotEmbedding.

#### Pseudotime ordering of cardiogenic trajectory

A pseudotime trajectory approximating the differentiation of progenitor cell types to mature cell types was curated using the addTrajectory function (preFilterQuantile = 0.9, postFilterQuantile = 0.9) to order cells along the trajectory backbone C6, C5, C12, C13, C7, C8, C14. This backbone represents the biologically relevant cardiogenic differentiation path; epiblast, EomesME, Mesp1ME, LPM2, LPM1, CMs/CPs. We leveraged ArchR’s series of pseudotime vector calculations to fit and align individual cells based on their Euclidean distances to the defined backbone’s cell type clusters’ mean coordinates in order to fit a continuous trajectory path in batch corrected LSI dimensional space. This resulting path with scaled, per-cell pseduotime values was then visualized in UMAP space using the plotTrajectory function. We then performed an integrative analysis to identify positive TF regulators along trajectory pseudotime. We integrated gene accessibility scores and gene expression data with motif accessibility across pseudotime using correlateTrajectories function and visualized correlated matrices in trajectory space with plotTrajectoryHeatmap function.

#### Assessment of positive transcription factor regulators

A putative positive regulator represents a TF whose gene expression is positively correlated to changes in accessibility of its corresponding motifs. Using the previously calculated motif z-score deviations, we stratified motif z-scores variation between all clusters to identify the maximum motif z-score delta. We next used the correlateMatrices function to correlate motifs to gene expression in batch-corrected LSI dimensional space, then used these correlations to identify motifs with maximized deviance from expected accessibility averages in other cells, and ranked TFs accordingly. We required positive TF regulators to have correlations greater than 0.5 (and adjusted p value < 0.01) between their gene expression and corresponding motifs, and deviation z-scores with maximum inter-cluster variation difference in the top quartile (quantile 0.75). Correlations were plotted for visualization using the ggplot function. While the ranking association with analysis might be vulnerable to generating false-negatives, wherein potential TF drivers aren’t recognized, we found overlay of motifs with TF gene expression and gene score values along the cardiogenic trajectory and in UMAP cluster space served to sufficiently identify the highest confidence drivers.

#### Differential peaks and differential motif enrichment comparisons between cell types

Pairwise comparisons between cell types of accessible peak differences was performed using the getMarkerFeatures function (Wilcoxon test, TSS enrichment and log10(nFrags) bias, 100 nearby cells for biased-matched background, 0.8 buffer ratio, 500 maximum cells) by setting one cell type as the lead comparison (useGroup) and one cell type as the relative comparison (bgdGroup). These pairwise comparisons of differential peaks were saved as .RDS objects and served as inputs to the (peak,gene) association analyses with rGreat (below in Methods). Differentially enriched peaks (FDR<= 0.05, abs(Log2FC) >=1) were visualized as MA plots. Motif enrichment of differential peaks was determined using the peakAnnoEnrichment function (FDR<= 0.05 and Log2FC >=1 for useGroup enrichment or else Log2FC<= -1 for bgdGroup enrichment) to determine motifs enriched in differential peaks between cell type groups. Enriched motifs were rank-sorted and colored by significance of enrichment, then plotted using the ggplot function.

#### Assessment of peak-to-gene linkages

Peak-to-gene linkage analysis to assess correlations between chromatin accessibility and gene expression was performed using the addPeak2GeneLinks function on batch corrected LSI dimensions (correlation cut off > 0.45, FDR < 1E-4, resolution 1000 bp for optimized browser track visualization). Peak-to-gene linkages for differentially expressed genes (identified in scRNA-seq analyses) were visualized with cell type cluster browser tracks using plotBrowserTrack.

### Association between scATAC-seq differential peaks and scRNA-seq differentially expressed genes

The rGREAT1(v1.26.0) bioconductor R package (Gu, 2022) was used to generate gene lists linked to scATAC-seq differential peaks based on gene regulatory domains defined as 5 kb upstream, 1 kb downstream of the Transcription Start Site (TSS) and up to 100 kb to the nearest gene. The log Fold Change (logFC) for the (peak,gene) pairs where the peak was differentially accessible (FDR <= 0.05, Log2FC >=1) were plotted to show how the log fold change of the gene expression is associated with the log fold change of the accessibility of peaks. The (peak,gene) pairs in the top-right quadrant (Q3) of the plot correspond to differentially open peaks linked with genes whose expressions are up-regulated. Similarly, the (peak,gene) pairs in the bottom-left quadrant (Q1) correspond to differentially closed peaks linked with genes whose expressions are down-regulated. Fisher’s test (Pearce, 1992) was performed on the counts of (peak, gene) pairs in each of the four quadrants; up-regulated genes:differentially open peak regions, down-regulated genes:differentially closed peak regions, up-regulated genes:differentially closed peak regions and down-regulated genes:differentially open peak regions. This provided an estimate of the ratio of the odds of upregulated genes linked to differentially open peak regions versus the odds of up-regulated genes linked to differentially closed peak regions.

### Whole mount fluorescent *in situ* hybridization experiments

Validation of spatial gene expression and differentially expressed genes was conducted in stage-matched, littermate whole-mount embryos. The assay for whole-mount embryo in situ was adapted from the optimized whole-mount zebrafish embryo protocol using the RNAscope Multiplex Fluorescent Reagent Kit v2 and ProteasePlus (ACDBio) for embryo permeabilization as previously described (Gross-Thebing, Paksa & Raz, 2014; Soysa et al., 2019). De-yolked whole embryos were fixed in 4% paraformaldehyde solution (Electron Microscopy Sciences 15710) overnight at 4°C. Embryos were then washed 2x in PBST and processed through 10 min incubations in a dehydration series of 25%, 50%, 75%, 100% methanol on ice. Embyros were stored in 100% methanol at −20°C short term until initiation of the *in situ* hybridization protocol. Yolk sac DNA or anterior proximal extraembryonic regions prior to fixation were used for genotyping. Catalogue numbers for ACDBio RNAscope probes used in this study: eGFP (400281-C1, -C2, -C4), Tdgf1 (506411-C1), Fgf8 (313411-C1), Eomes (429641-C2), Myl7 (584271-C3), Anxa2 (501011-C2), Nkx2-5 (428241-C2). Whole-mount embryos were imaged in cold PBS using an upright epifluorescent microscope (Leica MZFLIII, Leica DFC 3000G, Lumen Dynamics XCite 120LED) and acquisition software LASX (Leica). Control and *Mesp1* KO embryo comparisons were imaged and processed with identical parameters.

### Whole-mount embryo X-gal staining and imaging

X-gal staining for LacZ enhancer activity was performed according to standard protocols (Anderson et al., 2004; Materna et al., 2018; Sinha et al., 2015; Wilkinson & Nieto, 1993). Briefly, embryos were fixed in 4% paraformaldehyde at 4°C and stored in PBS until initiation of standard X-gal staining protocol. Littermate embryos were processed and imaged identically and simultaneously in brightfield using a Leica MZ165 FC stereomicroscope with DFC450 camera. Genotyping was done following blind processing.

### Whole-mount embryo immunostaining and light sheet imaging

Dissected embryos were fixed in 4% paraformaldehyde for 1 hour at room temperature with gentle agitation, washed in PBS, and stored in PBS + 0.2% sodium azide short-term at 4°C until initiation of immunostaining. Immunostaining was performed in PCR strip tubes. Embryos were incubated in blocking solution; PBS + 5% normal donkey serum, 0.2% sodium azide, 0.5% Triton X-100 (Sigma, X100-500 mL) with 100 μg/mL unconjugated Fab fragment donkey anti-mouse (Jackson Immunoresearch, 715-007-003) for 2 hours at 37°C with gentle rocking agitation. Following PBS washes, primary staining was done in blocking solution overnight and subsequently washed with PBS. Secondary staining incubation was done in blocking solution for 2-3 hours protected from light, and embryos were subjected to final PBS washes. All steps of immunostaining protocol were done at 37°C with gentle rocking and rotation. Antibodies used in this study: sheep polyclonal Foxc2 (R&D, AF6989), chicken polyclonal GFP (Aves, GFP-1020), rabbit polyclonal Cre (Millipore, 69050). Light sheet embryo images were acquired using Z1 Light Sheet Microscope (Zeiss) and processed as described (Dominguez et al., 2022).

## ACKNOWLEDGEMENTS

We thank members of the Bruneau Lab for thoughtful discussion, comments, and advice on the study, Kathryn Claiborn for editorial assistance, Junli Zhang for TARGATT transgenic mouse microinjections, M. Ryan Corces for advice on ArchR analysis, and Angelo Pelonero for general bioinformatics advice. We thank the UCSF Center for Advanced Technology for sequencing, and the Gladstone Institutes Bioinformatics, Genomics, and Microscopy and Histology cores for technical advice and support.

## AUTHOR CONTRIBUTIONS

A.L.K. and B.G.B. conceived and designed the study, and interpreted the data. S.A.B.W., A.L.K., and S.S.R. performed embryo dissections. S.A.B.W. managed animal husbandry and genotyping. S.A.B.W. and A.L.K. performed whole mount *in situ* hybridization experiments. A.L.K. performed imaging and image processing of embryos used for sequencing library preparations, and imaging and processing of *in situ* hybridization embryos. A.L.K. and S.S.R. generated scRNA-seq and scATAC-seq libraries. A.L.K. performed scRNA-seq analyses. A.L.K. and K.C. performed scATAC-seq analyses with input from S.S.R. and R.T. A.A. performed peak,gene association analysis with input from R.T. W.P.D. generated the *Hipp11^Smarcd3-F6::eGFP^* mouse and first observed the posterior *Smarcd3*-F6 phenotype in *Mesp1* KO embryos. T.S. generated, processed, and imaged *Tdgf1* enhancer transgene embryos. M.H.D. generated immunostaining and light sheet imaging embryo data. B.L.B. and D.S. supervised and advised. A.L.K. prepared figures and wrote the manuscript with input from co-authors.

## DATA AVAILABILITY

Raw and processed data for the whole embryo scRNA-seq and scATAC-seq datasets reported in this paper are available through the Gene Expression Omnibus (GEO) with accession code GSE210639. All analysis software is referenced in Methods section and is freely available from respective developers. Analysis scripts used to generate figure panels are freely available from the authors upon request.

